# Adding stochastic negative examples into machine learning improves molecular bioactivity prediction

**DOI:** 10.1101/2020.05.21.107748

**Authors:** Elena L. Cáceres, Nicholas C. Mew, Michael J. Keiser

## Abstract

Multitask deep neural networks learn to predict ligand-target binding by example, yet public pharmacological datasets are sparse, imbalanced, and approximate. We constructed two hold-out benchmarks to approximate temporal and drug-screening test scenarios whose characteristics differ from a random split of conventional training datasets. We developed a pharmacological dataset augmentation procedure, Stochastic Negative Addition (SNA), that randomly assigns untested molecule-target pairs as transient negative examples during training. Under the SNA procedure, ligand drug-screening benchmark performance increases from R^2^ = 0.1926 ± 0.0186 to 0.4269±0.0272 (121.7%). This gain was accompanied by a modest decrease in the temporal benchmark (13.42%). SNA increases in drug-screening performance were consistent for classification and regression tasks and outperformed scrambled controls. Our results highlight where data and feature uncertainty may be problematic, but also show how leveraging uncertainty into training improves predictions of drug-target relationships.

## INTRODUCTION

Machine learning and deep neural network (DNN) methods have made great strides in scientific pattern recognition, particularly for cheminformatics^1–7^. As larger amounts of training data (molecules and their protein binding partners) have become publicly available, ligand-based predictions of polypharmacology have expanded from classification of binding (e.g. active/inactive) to regression of drug-target affinity scores (e.g., K_i_, IC50)^3,4,8–12^. These models exploit the similar property principle of chemical informatics, which states that small molecules with similar structures are likely to exhibit similar biological properties, such as their binding to protein targets^13^. Such approaches assume that the principle holds true for large datasets and hinge on the expectation that a greater diversity of training examples will increase the likelihood of a model finding generalizable patterns relating chemical structure to bioactivity. However, these techniques may learn biased patterns from incomplete data for drug discovery and screening^14^. Academic cheminformatic machine learning training sets derive from public bioactivity databases such as ChEMBL and PubChem BioAssay (PCBA)^15,16^. Theoretically, the more researchers who deposit their data, the more diverse the database. However, as scientific literature is a major contributor to these databases, a publication bias toward molecules with positive binding profiles (Figure 1) could skew both the dataset and, consequently, the resulting machine learning models predictions, as in Kurczab, et al^17^. We explore the feasibility of a method that leverages uncertainty in unexplored chemical space to augment incomplete public data for small molecule activity prediction using deep learning for both classification and regression.

**Figure 1.**
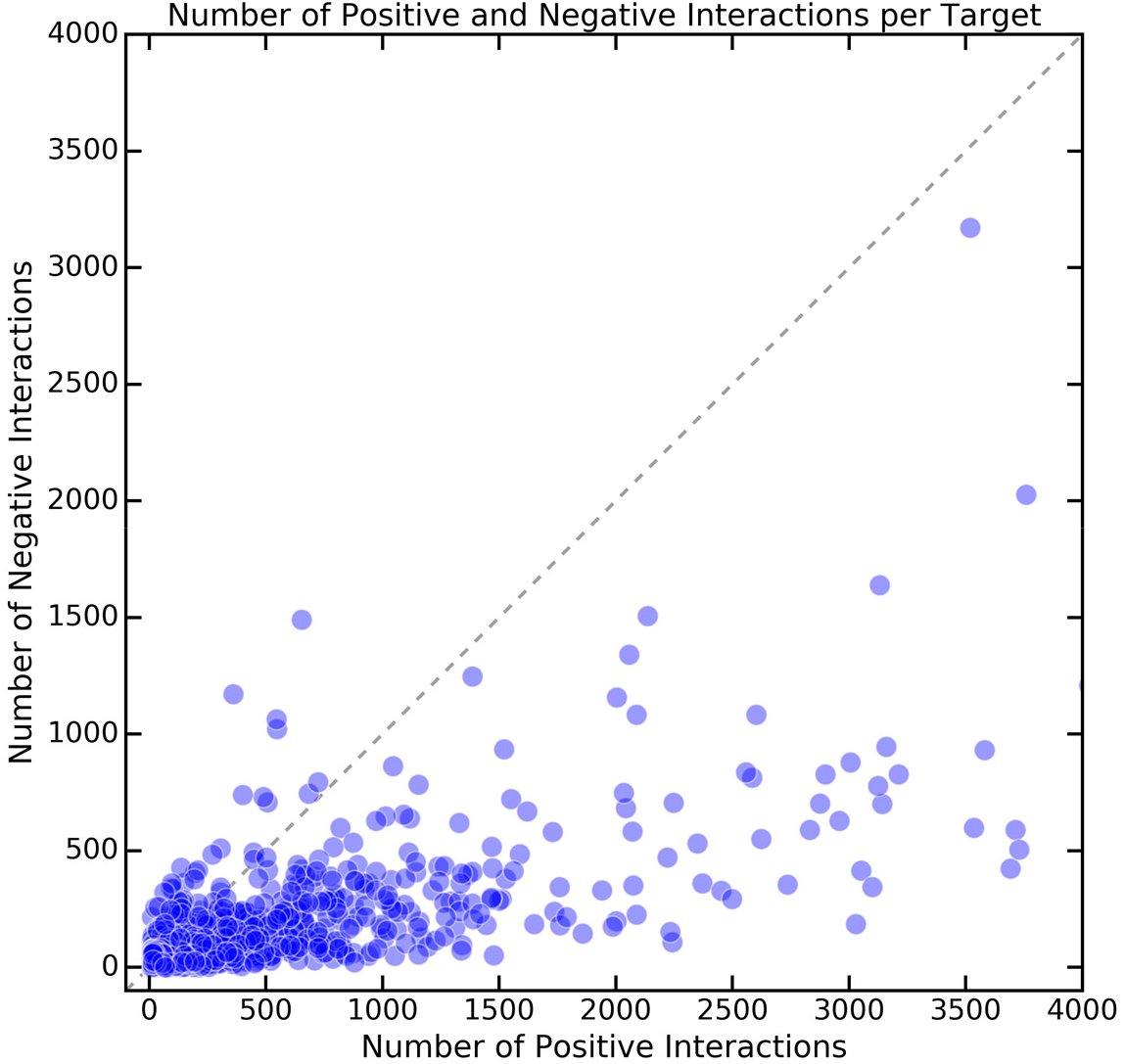
Protein targets are biased for positive interactions in a ChEMBL20 training set. Target-wise distribution of binding (ligand) vs non-binding molecules in the training set. Each point represents a single protein target drawn from ChEMBL20. A 1:1 ratio of binders (positives) and non-binders (negatives) would fall along the dotted line.

A substantial literature focuses on correcting the balance of positive to negative examples (here, binders to non-binders) in machine learning training datasets as well as addressing dataset sparsity^12,18–25^. These corrections primarily adopt majority- or minority-based approaches. Minority-based approaches oversample underrepresented classes, generally accomplished by upweighting or oversampling existing training examples or by adding similar synthetically-generated ones^19,24^. Majority-based approaches typically undersample the overrepresented class in order to achieve balance. Many class imbalance approaches address situations where positive examples were in the minority. This presents a unique problem for cheminformatic datasets where binders (<10uM) are frequently the majority class and non-binders are the minority reported class (Figure 1), despite binding in comprehensive screens being a rare event^26^. For cheminformatic datasets, undersampling the majority class members would minimize the crucial effort researchers have invested to establish the chemical feature diversity upon which similar property principle-based approaches rely^25^. Accordingly, De la Vega de Leon et al. found that removing/ignoring activity labels can decrease performance in proportion to the amount of data removed^22^. As non-binding molecules typically arise from the same series as binders, and consequently share many of their chemical features, we suspected that oversampling existing negative examples would contribute little to expanding a model’s decision boundaries. It would follow, therefore, that oversampling may fail to add diversity to the minority class, whereas methods that rely on synthetic interpolation (i.e. generating new fingerprints very similar to existing negatives) increase the chance of mislabeling a new ligand in the chemical series and overlook protein targets lacking negative pharmacology data^19^. From a machine learning perspective, this hinders a model’s generalizability and the scope of its chemical feature space. In turn, oversampling negative examples would seem especially problematic for cheminformatic datasets.

Random sampling of unassayed chemical space to assert weak but diverse negative examples may address these concerns. Others have shown that incorporating random negative data into training improves classification performance by SVMs^27^, potency-sensitive influence relevance voters^28^, and Bernoulli Naïve Bayes classifiers^29^. Kurczab et al. assessed the influence of negative data on a set of eight targets and found that a ratio of 9:1 to 10:1 of negatives to positives were favorable for classification^30^. In this work, we introduce putative negatives that continuously change throughout training, and extend this method beyond classification to regression tasks for thousands of protein targets at once. We evaluate prediction performance on screening and temporal benchmarks and search for optimal positive-to-negative ratios under both test scenarios.

We propose an online (continuous) pharmacological training augmentation procedure for regression and classification tasks: stochastically oversampling the minority (non-binder) class from the pool of unlabeled molecule-to-protein interactions spanning the molecule vs protein target training space. We designed Stochastic Negative Addition (SNA) with the challenges of ligand-based drug design in mind. SNA adds more molecule-protein pairs to a training set where negative examples are otherwise outnumbered and/or unevenly distributed. Paradoxically, whereas most molecules do not bind to most proteins, the literature-based pharmacological datasets we used contain a preponderance of positive reports (Figure 1): We intended SNA to counter this trend without overwhelming training with negative examples. This method encodes uncertainty for unstudied, and unlabeled, drug-target pairs. It exploits the observation that, despite meaningful cases of unexpected polypharmacology, ligand binding events at ≤10 uM are comparatively rare^26^. This study expands on prior work by investigating the effect of training augmentation for large numbers of protein targets in a multitask setting, applying the method to regression tasks, and assessing the impact of random negatives on complementary benchmarks.

We assessed DNN model performance on two external test sets: First, we created a Time Split hold-out to address a drug discovery scenario with the understanding that this test set would be skewed toward having fewer negatives. Second, we created a complementary “screening” use-case benchmark, with a preponderance of negatives. For it, we used the densely assayed Drug Matrix collection^31,32^, and removed all of its protein-molecule interactions that intersected with the training set to avoid data leakage. To determine how much pre-existing negative examples contributed to performance, we trained alternative DNNs where we removed negatives from the training dataset. We explored whether SNA could rescue performance in this scenario where actual negatives were absent. We compared these models to an unaugmented, standard training regime and appropriate adversarial control studies^33^. We then explored whether different ratios of binders to non-binders affected performance. Finally, we evaluate whether SNA improves classification to the extent that it does regression. We find using *SNA* with a one-to-one positive-to-negative ratio improves performance on screening scenarios with minor penalty to temporal benchmarks.

## RESULTS

### Adding stochastic negatives improves regression performance

We posited that existing sparse public datasets omit much of the chemical diversity of negative bioactivity space. To address this, we developed a machine learning training procedure to transiently add likely negative examples: unstudied pairs of small molecules and protein targets that we assert to not bind. Using a Stochastic Negative Addition (SNA) procedure, model predictions on a screening scenario benchmark dataset (Drug Matrix) improved with minimal loss to performance on a temporal test benchmark (Time Split) (Figure 2).

**Figure 2.**
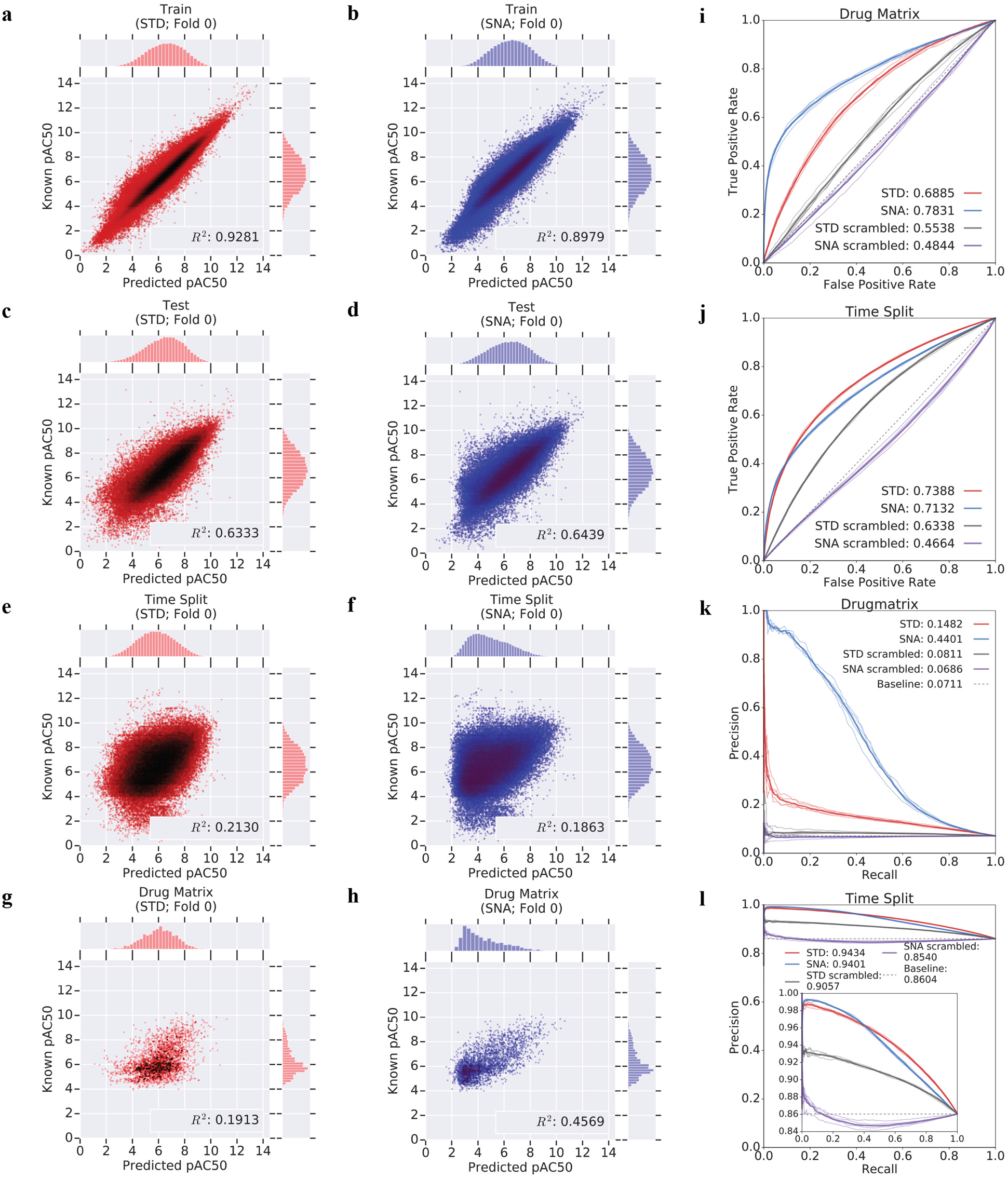
SNA markedly improves screening performance at minimal cost to temporal benchmarks. Predictions from a regression-based deep neural network model trained on available data (*STD;* red; a,c,e,g) and the same model trained with the addition of stochastically-chosen negative examples during each epoch (*SNA*; blue; b,d,f,h) show R^2^ improvements on Drug Matrix (e,f) with minimal cost to Time Split (g,h) performance. Drug Matrix screening benchmark improvements translate to regression-as-classification AUROC_r_ (i) and AUPRC_r_ (k) plots where the *SNA* DNN (blue) outperforms the *STD* DNN (red). Time Split AUROC_r_ (j) and AUPRC_r_ (l) plots show small decreases to performance on temporal hold out predictions from *SNA* models (blue). Performance on both hold-out datasets outperform scrambled controls (*STD scrambled* - grey; *SNA scrambled* - purple), with SNA models showing greater gains over their random counterparts. *SNA scrambled* models (i-l; purple) move the baselines for regression-as-classification closer to 0.5 for AUROC_r_ and the positive-to-negative ratio (dotted line) for AUPRC_r_.

DNN models trained with five-fold cross validation using SNA (hereafter denoted in italics as *SNA*) outperformed conventionally-trained models (standard; *STD*) on the screening (Drug Matrix) benchmark (Figure 2(g,h); Table 1; Supplementary Table 1; Supplementary Figures 9-10, 17-18) with little effect on training or random test performance (Figure 2(a-d); Table 1; Supplementary Table 1; Supplementary Figures 9-10, 17-18). *SNA* performance increased 122% in R^2^ over the STD model on Drug Matrix affinity pAC_50_ values (see Methods). As with most screens, much of the data within the screening benchmark consisted of first-pass “primary” observations assessed only at the single dose of 10 μM. Regression could not be performed on these observations, as no dose-response curve had been collected. To assess the effect of the proposed SNA training procedure on classification tasks, which would include these cases as well, we used two analyses: classification and regression-as-classification. The former consisted of training equivalent DNN architectures with classification loss functions -- see dedicated section below. For the latter, we evaluated the output of the original regression models as classifiers *post hoc*, by thresholding affinity into positive and negative assignments according to pAC_50_ for the underlying truth values and constructing AUPRC_r_ and AUROC_r_ metrics over a range of affinity thresholds instead of confidence thresholds. Thus, we calculated regression-as-classification AUPRC_r_ and AUROC_r_ by combining primary negatives with secondary (dose-response) negatives from the Drug Matrix screen versus secondary positives (see Methods). This analysis on Drug Matrix showed a 196% increase in AUPRC, and 14% increase in AUROC_r_ for models trained using *SNA* over *STD* (Figure 2(i,k); Table 1; Supplementary Table 1; Supplementary Figures 9-10, 17-18).

**Table 1.**
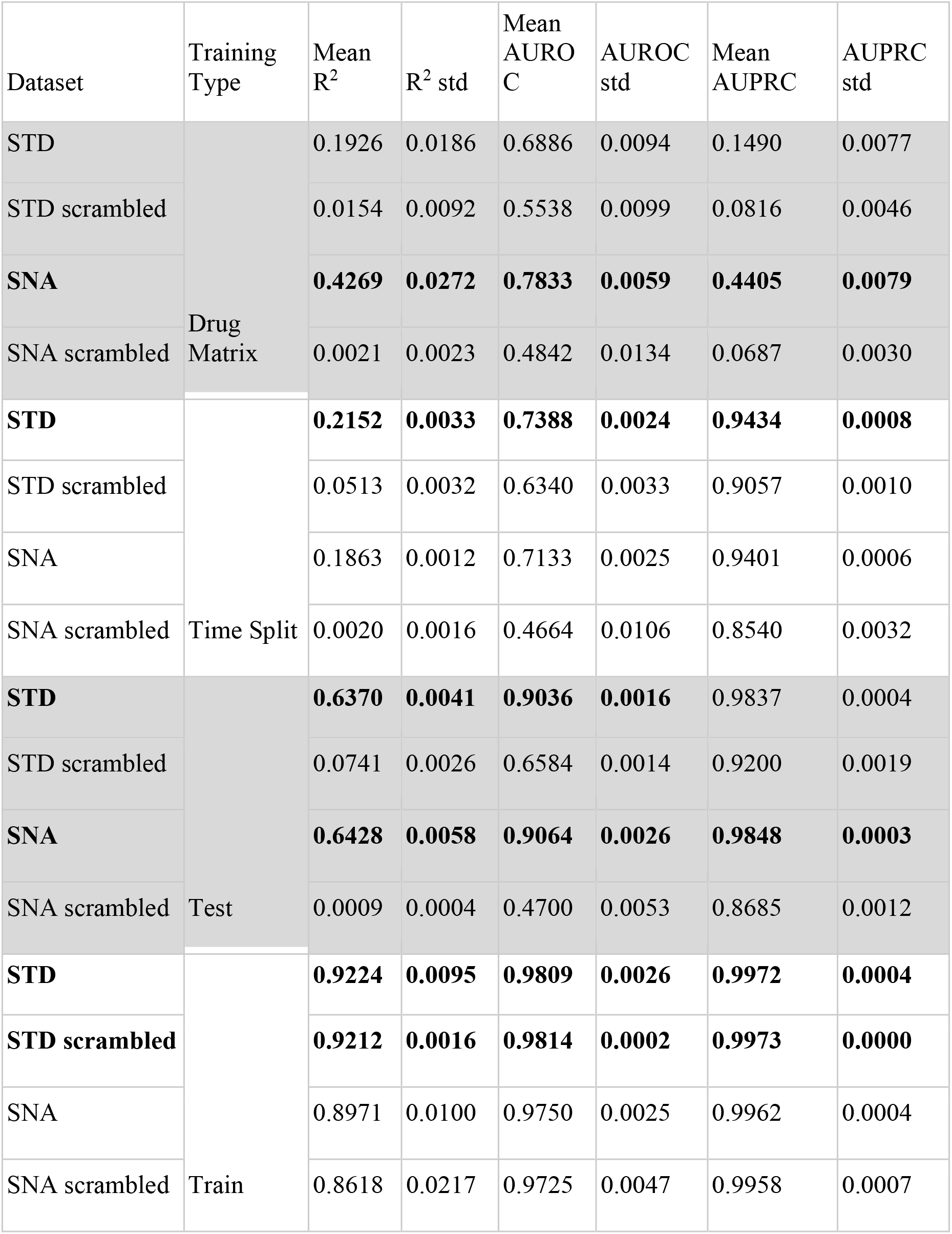
Mean and standard deviation (std) across independent 5-fold cross validation for *STD* and *SNA* DNN models.

By contrast, *SNA* performance on the temporal (Time Split) benchmark decreased slightly, with *SNA* models decreasing by 13% in R^2^ and 3% in AUPRC_r_ compared to *STD* (Figure 2 (e-f,l); Table 1; Supplementary Table 1; Supplementary Figures 9-10, 17). *STD* and *SNA* models generalized similarly on cross-validation test sets (Figure 2(c-d), Table 1; Supplementary Table 1; Supplementary Figures 9-10, 17-18), whereas standard models more precisely recapitulated their exact training examples ((Figure 2(a-b)), Table 1; Supplementary Table 1; Supplementary Figures 9-10, 17-18) than the equivalent *SNA* model, as expected.

### SNA brings scrambled control models closer to theoretical random for regression

To evaluate whether the models withstood adversarial controls^33^, we trained models on molecules whose annotations to protein targets had been randomized (y-randomization)^34–36^. *SNA scrambled* and *STD scrambled* models were trained with and without SNA procedures, respectively. Our goal was to verify that these intentionally scrambled models would underperform equivalent non-scrambled models on actual benchmarks. Thus, as in previous sections, we evaluated these models on screening, temporal, and 5-fold cross validation (Test) sets.

As intended, scrambled models greatly underperformed those trained on actual data (Figure 3; Table 1; Supplementary Table 1; Supplementary Figures 17-18). However, some empirically-scrambled models using standard training exceeded expected theoretical performance for balanced models (Figure 3 (e-h); Table 1; Supplementary Table 1; Supplementary Figures 17-18). Scrambled models converged during training and achieved high performance on their scrambled train datasets (Supplemental Table 1; Supplementary Figures 13-14, 17-18), consistent with potential dataset memorization rather than generalization^37^. Unsurprisingly, the R^2^ for scrambled models neared 0.0 for screening, temporal, and cross-validation sets (Supplementary Figures 13-16). While models trained on actual data outperform their scrambled controls, these controls exceeded frequently used, theoretical baselines such as 0.5 for AUROC_r_ and the positive-to-negative ratio random baseline for AUPRC_r_ in regression-as-classification analyses. *STD scrambled* models outperformed the 0.5 theoretical-random in AUROC_r_ (Drug Matrix screening set: 0.5538±0.0099; Time Split temporal set: 0.6340±0.0033) (Figure 3(e,f); Supplementary Table 1). We also found these *STD scrambled* models performed better than the random prevalence line in AUPRC_r_ for Drug Matrix (random prevalence: 0.0711; AUPRC_r_: 0.0816±0.0046) and temporal benchmarks (random prevalence: 0.8604; AUPRC_r_: 0.9057±0.0010) (Figure 3 (g,h); Supplementary Table 1). *SNA scrambled* models exhibited AUROC_r_ and AUPRC_r_ nearer the expected theoretical random baselines in both the Drug Matrix benchmark (AUROC_r_: 0.4842±0.0134, AUPRC_r_: 0.0687±0.003) and temporal benchmark (AUROC_r_: 0.4664±0.0106, AUPRC_r_: 0.8540±0.0032) (Figure 3; Supplementary Table 1).

**Figure 3.**
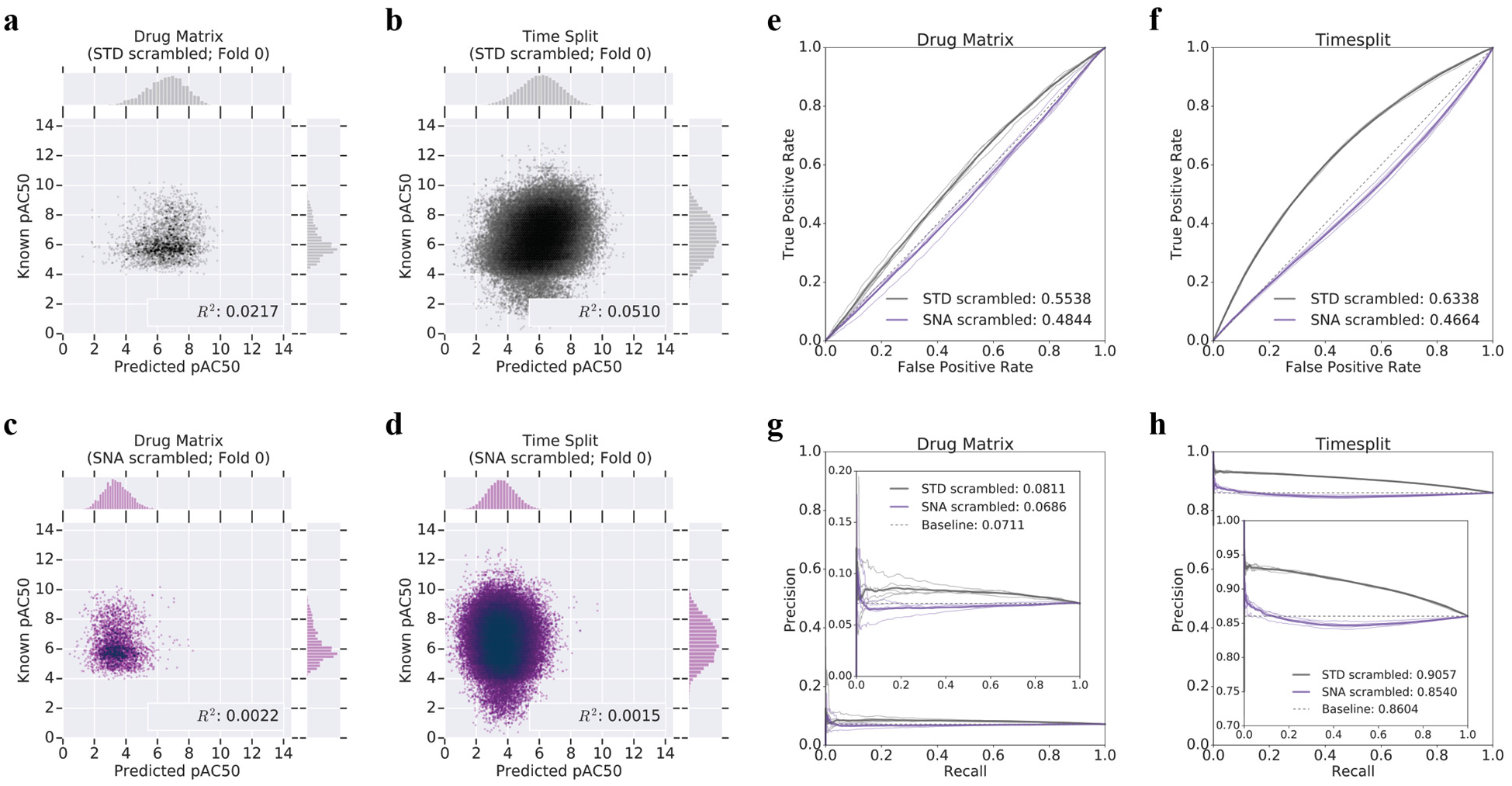
Empirically scrambled control models using SNA more closely match expected random baselines. R^2^ plots of a representative fold from 5-fold cross validation (a-d) illustrate the predictive behavior of unmodified DNN models trained with y-randomized data (*STD scrambled;* grey; a,b) and the equivalent networks trained with the stochastic negative procedure (*SNA scrambled*; purple; c,d). Baselines for regression-as-classification AUROC_r_ (e,f) show the *SNA scrambled* random line (purple) is closer to 0.5 (dotted line) than *STD scrambled* (grey). This is true for both Drug Matrix (e) and Time Split (f). Regression-as-classification for AUPRC_r_ (g, h) shows a similar trend where *SNA scrambled* baselines (purple) approach the ratio of positives-to-negatives in the Time Split hold out (dotted line) especially when compared to *STD scrambled*(grey).

### SNA improves performance for classification models

To evaluate whether the SNA training procedure was stable beyond regression and regression-as-classification uses, we developed and evaluated DNN classifiers with similar architectures. As with regression models, *SNA* classifiers saw increased model performance for the Drug Matrix screening benchmark, with a minor decline on the TimeSplit temporal benchmark (Supplementary Table 2; Figure 4; Supplementary Figures 17-18). In 5-fold cross-validation, *SNA* improved screening benchmark performance by 151% AUPRC and 13% AUROC (Supplementary Table 2; Supplementary Figures 17-18). Consistent with regression models, classification networks trained with *SNA* exhibited minor (4% AUROC and a 1% AUPRC) decreases on the Time Split benchmark (Supplementary Table 2, Supplementary Figures 17-18). As before, both models outperformed their scrambled baselines. Classifier DNNs showed less performance gain over random controls in the temporal benchmark than regressor DNNs (Figure 4 (e,f)).

**Figure 4.**
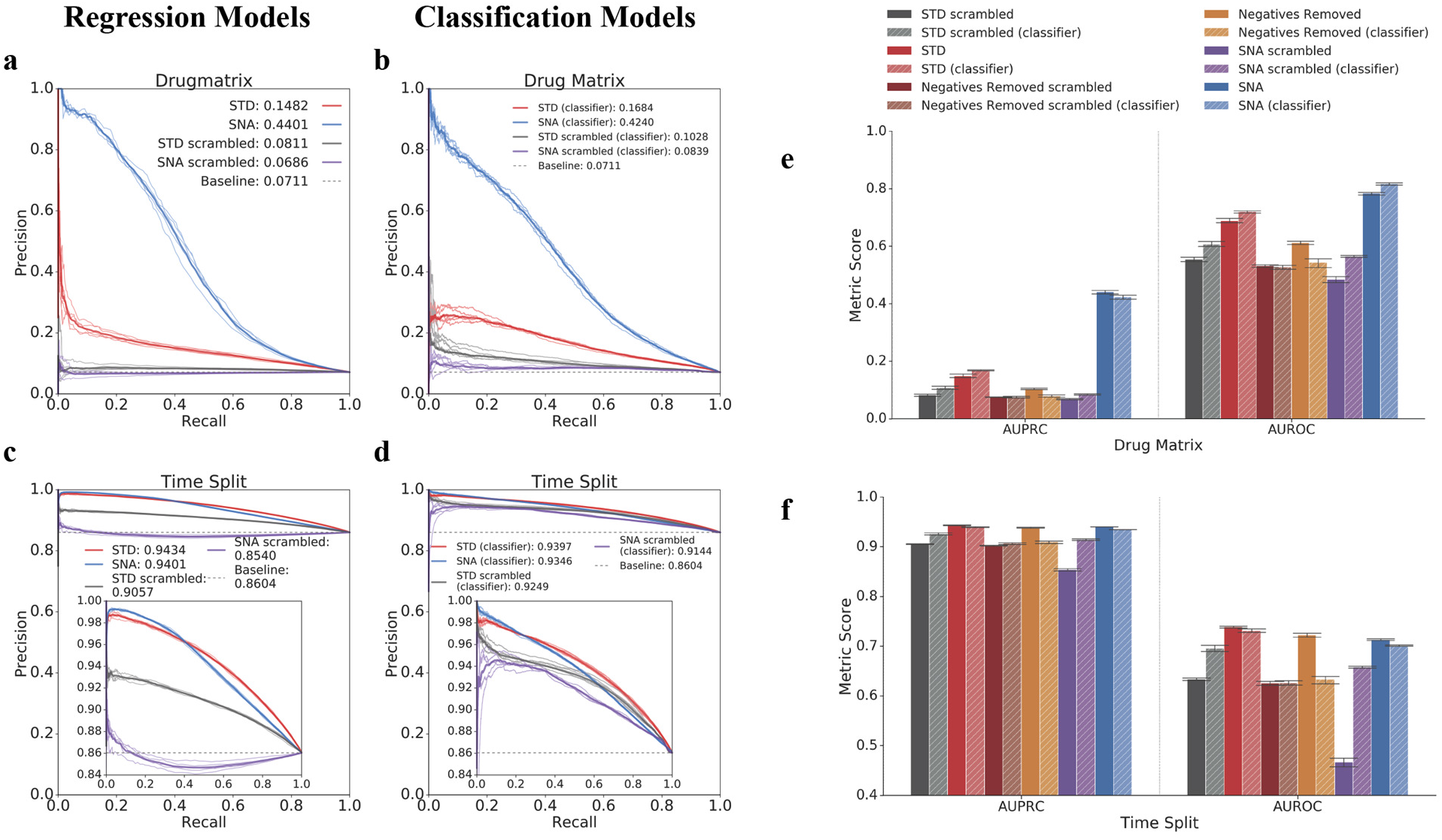
Classification DNN models show similar, but not identical trends to regression-as-classification evaluation on regression models. AUPRC on 5-fold cross-validated classification networks (b,d,e,f) versus equivalent regression networks (a,c,e,f). For both classification (a,e) and regression tasks (b,e), networks trained with SNA (*SNA*; blue) improved Drug Matrix AUPRC compared to DNNs trained without SNA or other data augmentation (*STD*; red). However, *SNA* models (blue; solid - regression, hashed - classification; c,d) did not improve or even decreased Time Split performance compared to *STD* (red; solid - regression, lightened and hashed - classification; c,d). AUROC and AUPRC bar plots for Drug Matrix (e) and Time Split (f) illustrate differences between classification (lighter, hashed bars) and regression models (solid bars) for SNA models (blue, green), their equivalent networks without SNA (red, orange), and y-randomized controls (grey, purple, sienna, chartreuse). All DNN models but the *Negatives Removed* classification model (e,f; light orange; hashed) outperformed the scrambled benchmarks (e,f; sienna; hashed). Regression *STD* models (red; solid) underperformed classification *STD* models (light red; hashed) for Drug Matrix (e) and Time Split (f), but regression *Negatives Removed* models (orange; solid) outperformed classification *Negatives Removed* models for the same test cases. Regression (solid) models for *SNA* (blue) outperformed classification (light blue, hashed) AUROC and AUPRC for Time Split (f) and AUPRC for Drug Matrix (e), but underperformed for Drug Matrix AUROC.

### SNA improves regression models trained without negatives

As SNA improved performance on a training set where negatives are not guaranteed to be distributed across the benchmark sets in the same manner as the train set, we were curious whether SNA would improve cases where there are no true negative training data for ligand-binding prediction. To address this question, we evaluated two training regimes. First, we trained a DNN model solely on positive ligand-target examples (*Negatives Removed*). Second, we trained the equivalent *Negatives Removed* model, corrected by the SNA procedure (*Negatives Removed* +*SNA*). To compare model performance, we maintained the same benchmarks as before (screening/Drug Matrix, temporal/Time Split, and cross-validation). We hypothesized that removing all training negatives would damage model performance across the board, while incorporating SNA might partially rescue this effect. Additionally, we hypothesized classification models would be more sensitive to the removal of training negatives than regression models.

Broadly, regression models trained without negative examples underperformed by regression-as-classification metrics, while achieving similar or better R^2^ to standard (*STD*) for Drug Matrix and Time Split (Figure 5; Supplementary Table 1, 2). The R^2^ difference between *Negatives Removed* and *STD* models was minimal for the Drug Matrix screening benchmark (*Negatives Removed* R^2^: 0.1973±0.0176; STD R^2^: 0.1926±0.0186) (Figure 5(a); Figure 2 (g); Supplementary Table 1, Supplementary Figures 9, 11). However, we observed larger differences in AUROC_r_ and AUPRC_r_ where the STD model outperformed the equivalent *Negatives Removed* model for Drug Matrix (*Negatives Removed* AUROC_r_: 0.6120±0.0076 vs *STD* 0.6886±0.0094; *Negatives Removed* AUPRC_r_: 0.1039±0.0025 vs *STD* AUPRC_r_: 0.1490±0.0077) (Figure 5 e; Supplementary Table 1; Supplementary Figures 23-24). Removal of negatives from training harmed the Time Split temporal benchmark performance (−2.2% AUROC_r_ and −0.5% AUPRC_r_ change from *STD*) (Figure 3f; Supplementary Table 1), but these models showed minor improvements in R^2^ (9.3% increase from *STD* models) (Figure 5 (c,f)). For cross-validation (Test) and training-data benchmarks, removal of negatives during training uniformly decreased performance, by 5% (Test) and 15% (Train) in R^2^ (Supplementary Table 1).

**Figure 5.**
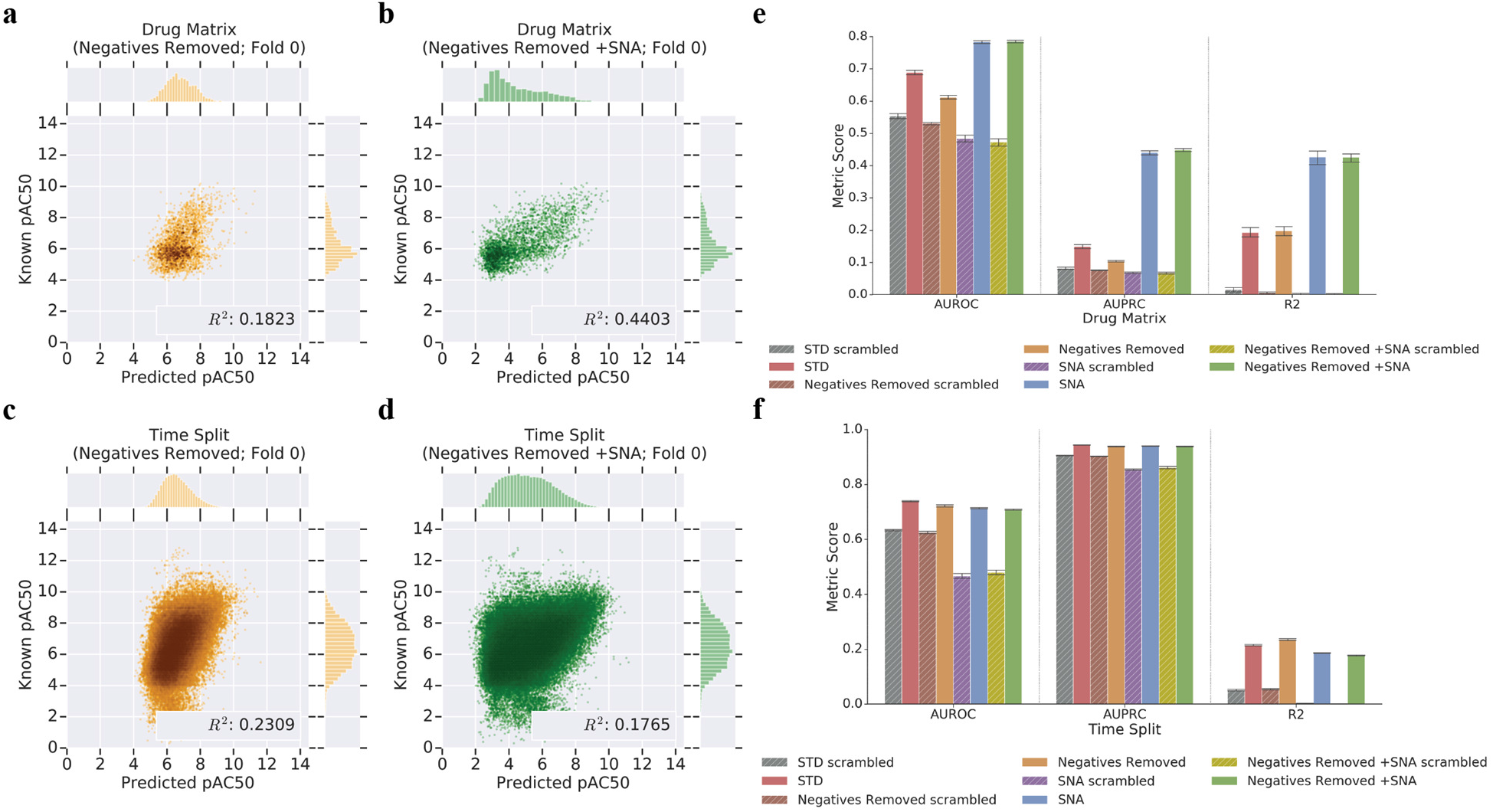
The SNA procedure rescues models trained without negative data. R^2^ plots (a-e) of the same fold from 5-fold cross validation show *Negatives Removed +SNA* models (green; b, d) rescue Drug Matrix predictive capability. Models trained with SNA in place of actual negatives suffer under the Time Split benchmark when compared to a model trained without negatives or SNA (Negatives Removed; gold; a, c). This Drug Matrix performance boost is consistent across regression-as-classification AUPRC_r_ and AUROC_r_ metrics on k-fold cross validated networks (e). For Drug Matrix, SNA models (e; blue, green) outperform equivalent DNNs with no stochastic negatives added (e; red, orange). SNA does not improve Time Split performance (f; blue, green) when actual negatives are lacking (f; red, orange). Scrambled controls for each experiment (e,f; grey, brown, purple, chartreuse) establish a baseline for performance. R^2^ metrics for *SNA scrambled* and *Negatives Removed +SNA* models are near zero for Time Split and Drug Matrix (e,f; purple, chartreuse).

We had anticipated that the SNA training procedure would partially mitigate the absence of true negatives during model training. Surprisingly, the *Negatives Removed +SNA* procedure yielded models with performance nearly indistinguishable from SNA models trained with full data, *SNA* (Figure 5 (e,f); Supplementary Table 1; Supplementary Figures 1-4). As with *STD* compared to *SNA*, *Negatives Removed +SNA* substantially improved the Drug Matrix screening benchmark performance while slightly decreasing that of the Time Split benchmark compared to a model trained with *Negatives Removed* alone. We observed 28%, 331%, and 115% increases to AUROC_r_, AUPRC_r_, and R^2^, respectively, for the Drug Matrix screening benchmark by adding SNA training to *Negatives Removed* models (Figure 5; Supplementary Table 1; Supplementary Figure 1). By contrast, we observed that the *Negatives Removed +SNA* model training decreased temporal benchmark R^2^ performance 24% to 0.1774±0.0018 compared to the *Negatives Removed* alone (Figure 5 (d,f); Supplementary Table 1, Supplementary Figure 2). We found little to no change in mean AUC metrics for regression of *Negatives Removed* or *Negatives Removed +SNA* models, suggesting that neither stochastic nor true negatives improve performance on the temporal benchmark. Under these performance metrics, the impact of stochastic negative data on model training could not be distinguished from that of true negatives. However, we did not find that stochastic negatives yielded any *greater* performance than true negatives, despite the greater diversity of chemical examples covered by the former. To address whether there was an advantage from reported negatives, we performed an alternative training analysis wherein we upweighted existing negatives during training to reach parity between positives and negatives (Supplementary Methods: Negatives Upweighted; Supplementary Table 1) and found little improvement to the Time Split benchmark.

### SNA training corrects for absence of true negatives in classification nearly as well as in regression

As with the regression models, removal of true negatives when training classification models damaged performance on most benchmarks. SNA predominantly rescued performance for classification *Negatives Removed* models. The removal of true negatives from classification DNN training so adversely impacted performance on holdout benchmarks that these models failed to exceed random baselines (Supplementary Table 2, Supplementary Figures 21-22). This was consistent with the expectation that classification models trained solely on positive data would overwhelmingly predict positive outcomes. Therefore, we expected that incorporating stochastically-imputed negatives during training (*Negatives Removed +SNA*) would improve classification. Drug Matrix screening benchmark performance improved markedly for *Negatives Removed +SNA* training compared to *Negatives Removed* models (48% increase in AUROC; 291% in AUPRC) (Supplementary Table 2, Supplementary Figures 21-22). *Negatives Removed* +*SNA* only slightly improved Time Split AUROC and AUPRC (3% to AUROC and 1% to AUPRC), although this was in contrast to regression models, where SNA had *decreased* performance in this scenario (Supplementary Tables 1,2, Supplementary Figures 21-22). Overall, we observed that the *Negatives Removed* regression model and its derived regression-as-classification interpretation outperformed the classification model on the screening benchmark. This was true as well for *Negatives Removed +SNA* training with the exception of AUROC_r_ for Drug Matrix.

### Restricting SNA by molecular similarity does not dramatically improve the procedure

To decrease the likelihood that SNA may assign true-but-unreported ligands to be negatives during training, we blacklisted potential molecule-target pairs by a separate cheminformatic method. This blacklist was created using the Similarity Ensemble Approach (SEA) to predict likely binders (Supplementary Information). We assessed the networks trained with the SEA blacklist (*SNA +SEA blacklist*) similarly to the base *SNA* model procedure for both classification and regression. As with *SNA* networks, the *SNA +SEA blacklist* DNNs outperformed *STD* models on Drug Matrix with minor decreases to Time Split for regression (Supplementary Table 1; Supplementary Figures 1-2) and classification (Supplementary Table 2; Supplementary Figures 5-6). The performance differences between *SNA* and *SNA +SEA blacklist* were minimal; typically within a 1% difference (Supplementary Table 1; Supplementary Figures 1-4). The exceptions were AUPRC_r_ and R^2^ where *SNA +SEA blacklist* outperformed *SNA* on Drug Matrix by 2.8% and 3.3%, respectively. The same was true for classification networks, with *SNA +SEA blacklist* performing within a 1% difference to *SNA* for all but AUPRC, where *SNA +SEA blacklist* induced an increase in the mean across cross validated models of 3% (Supplementary Table 2, Supplementary Figures 5-8).

### The optimal SNA ratio for DNN performance centers on 1:1 positive:negative examples

To assess the impact of the class balance ratio chosen for the SNA training procedure, we trained 14 networks with SNA minimum-ratios (i.e. minimum ratio of negatives-to-positives per protein target, below which negatives are added until the ratio is achieved) extending from no negatives added to the training (0% added) to 93%, as assessed on each protein target represented within a minibatch. We applied this procedure to regression and classification DNNs trained with *STD* and *Negatives Removed* contexts.

We found that the region between 40% to 60% added-negative ratio was the best tradeoff of performance across all benchmarks (Figure 6, Supplementary Tables 5,6). Consistent with established class-balance training procedures, a 50% or 1:1 addition of SNA appears ideal, for both classification and regression scenarios. We note that the Drug Matrix screening benchmark improvement is steepest between 10% and 30% negative-addition; while the Time Split benchmark suffers some decreases in this regime, they are far less pronounced than the improvements to the screening benchmark.

**Figure 6.**
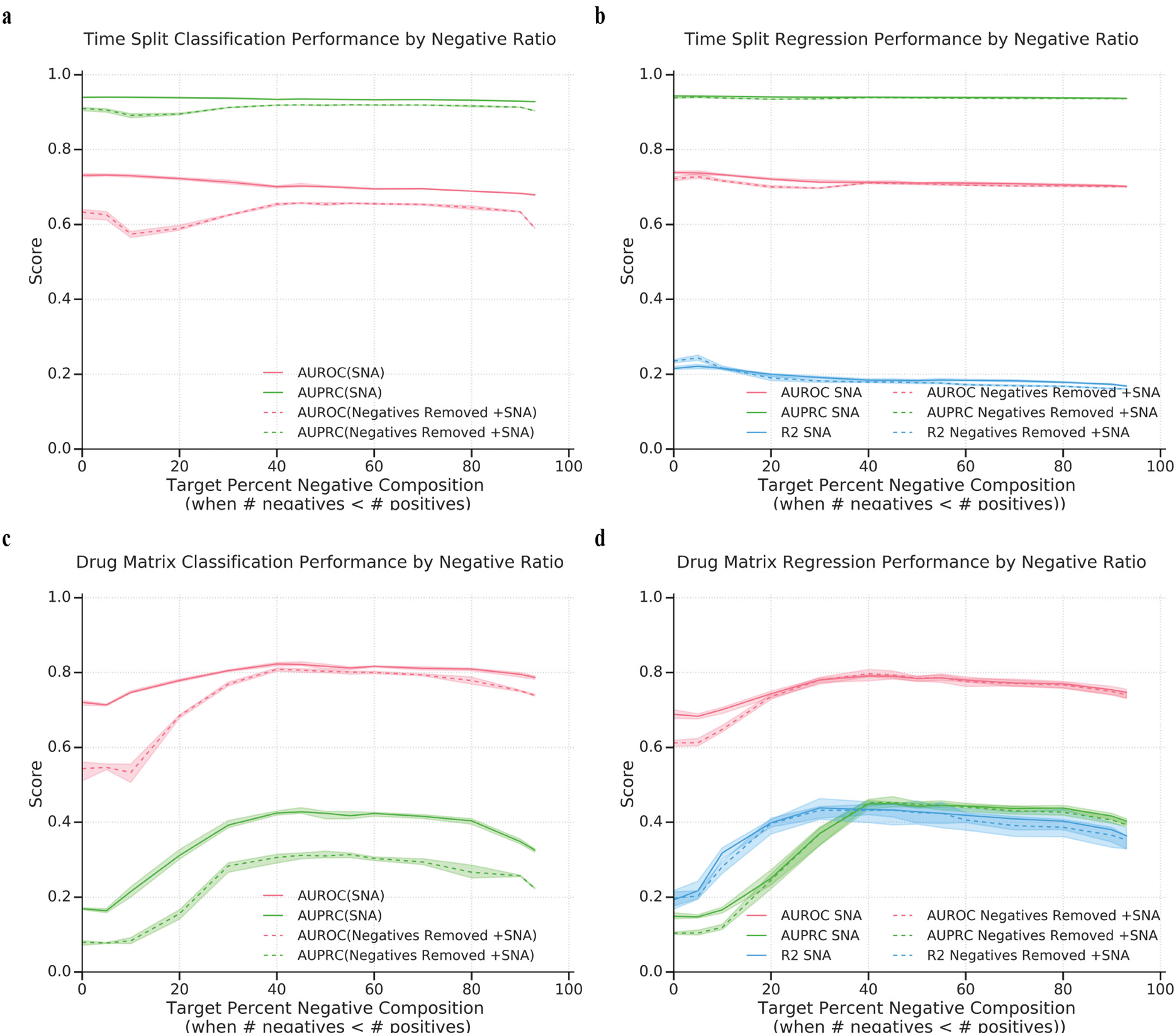
A balanced ratio of positives to negatives achieves the best overall performance. Increasing the target negative-to-positive ratio improves Drug Matrix performance for classification (c) and regression (d) up to ~40-50% negatives per target. This increase is observed with modestly decreased Time Split performance for classification (a) and regression (b) *SNA* models. Addition of negatives to a training set with negatives removed (*Negatives Removed +SNA;* dashed line) shows minimal effect on Time Split (a,b) and Drug Matrix (c,d) performance compared to the *SNA* model without negatives removed from the training set (solid line). Regression *Negatives Removed +SNA* models perform similarly to regression *SNA* DNNs (b,d). However, classification *Negatives Removed +SNA* networks perform worse than classification *SNA* models that were trained with pre-existing negative examples (a,c). Shaded areas represent the maximum and minimum boundaries within 5-fold cross validation.

The most exaggerated difference between classification and regression occurred for *Negatives Removed* models (Figure 6, Supplementary Tables 5,6). Regression models trained using the *Negatives Removed +SNA* method almost entirely rescued the all-data *SNA* model performance by a 40% negative-addition ratio for Drug Matrix (Figure 6 d). However, classification *Negatives Removed +SNA* could not match the AUPRC Drug Matrix screening performance of *STD* (Figure 6 c). The screening benchmark performance difference between *SNA* and *Negatives Removed* +*SNA* models was less pronounced by AUROC. For the Time Split benchmark, stochastic negatives were less effective at closing the performance gap between *Negatives Removed* and *STD* classification models (Figure 6 a) than they were for regression models (Figure 6 b). From this, we conclude classification tasks perform much better when trained on true negatives than when trained with stochastically imputed negatives. Regression models appear to see less gain from reported negatives as compared to stochastically imputed ones. However, as both classification and regression performed better with the addition of random negatives, we believe *SNA* can productively augment true negatives in the case when there are insufficient negatives and in the case when there are no negatives to speak of in the training set.

## DISCUSSION

In this study, we set out to investigate the impact of stochastic negative addition to DNN training. We evaluate DNN regression performance on small-molecule-to-protein target affinities, as well as thresholding the regressed values analogously to classification networks to obtain a regression-based AUROC_r_ and AUPRC_r_ performance metrics, which frequently agree with conventional classification task training and performance trends. We find that adding stochastic negatives to DNN training improves predictive performance on a full screening (Drug Matrix) hold-out benchmark for both classification and regression networks. We observe that this performance boost has minimal impact on a temporal evaluation scenario. We compare the results to scrambled baselines, which suggest the method does not solely rely on memorization for performance improvement.

Experimental screens and studies often focus on novel binders, optimizing for sensitivity and precision. Indeed, the literature-derived annotations mined from ChEMBL skew the training set toward positive examples, with 73% representing binding affinities at 10 uM or lower, 55% of 1 uM or lower, and 34% of 100 nM or lower. The remaining 27% of the training examples are explicit negatives—molecules that failed to inhibit the tested protein target by at least 50% at 10 uM. A surprising 6% of targets (138 proteins) have zero reported non-binders weaker than 10 uM. For training purposes, no targets have zero reported binders weaker than 10uM as outlined in Methods. We believe the dearth of negatives in training data derived from studies incentivized to publish positive results could contribute to poor machine learning model precision, such that false positives could flood the potential testing space in some cases. With this in mind, we hypothesized that adding transient random negative examples during training would improve model precision more than it would degrade model sensitivity. We find SNA improves the rate of negative assignments in exchange for a minor hit to performance on temporal splits (Figure 2). We consider this to be an acceptable trade-off in most applications, as false positives dilute search spaces for screening follow-up and manual review of promising binding candidates.

Ultimately, the comparative value of different test sets and measurement statistics are use-case specific choices that must be set by the researcher. The Time Split and Drug Matrix datasets represent two sides of a similar problem. Time Split attempts to approximate challenging and desired prospective validation performance across a broad range of models, where chemical novelty is foremost. As comprehensive negative data are so rarely publicly available, Drug Matrix represents a complementary scenario where a full matrix of molecules and proteins are provided which incidentally results in a higher ratio of negatives in the evaluation set. Inclusion of the Drug Matrix dataset is an attempt to quantify performance on highly imbalanced but realistic testing scenarios.

The SNA data augmentation procedure improves both regression and classification deep neural network performance when compared to a standard model (*STD*) (Figure 4, Supplementary Tables 1-2). Here, we define success as better performance on a screening-scenario benchmark, Drug Matrix, with a small relative hit to the Time Split benchmark. We find a balanced training dataset of positive and negative examples performs well (Figure 6, Supplementary Tables 5-6). The performance improvement consistently exceeded scrambled benchmarks, consistent with learning beyond simple memorization for each model.

While we find *SNA* improves Drug Matrix at a cost to Time Split performance, we note that the models contain negatives within the training set which may be unevenly distributed across tasks. This distribution may artificially boost performance for certain tasks with additional negative data. To address this, we train a model in the absence of negative data and without stochastic negatives as a sanity check where we expect reduced performance due to loss of relevant information. We find this *Negatives Removed* model improves generalizability for Time Split and depletes performance on Drug Matrix. One potential explanation for this performance gain may be a positive-reporting bias for novel molecules in the ChEMBL database underlying the temporal-split benchmark (71% positive, n=116929, Supplementary Table 4), which is derived from the literature and could reward models that are more likely to predict positive binding activity for a novel molecule, although we have not attempted to test this theory. We find *Negatives Removed* classification DNNs (Supplementary Table 2, Supplementary Figures 21-22) perform worse than *Negatives Removed* regression DNNs (Figure 5 e,f; Supplementary Table 1, Supplementary Figures 23-24). We hypothesize this may be due to regression models learning continuous relationships between chemical structure and bioactivity that may extrapolate into low-affinity regimes, whereas classification models entirely lack the negative categorical data and features around which to construct a binary decision boundary

Regression models trained with *Negatives Removed* exhibit performance losses which SNA can rescue (Figure 5). These data suggest that stochastic negatives may usefully supplement true negative data, but due to lack of clearly better performance; we do not believe that SNA should be used to supplant the use of true negatives in model training. SNA failed to completely rescue classification *Negatives Removed* performance. This may reflect fundamental differences between the aim of regression versus classification or the forms of the loss functions in question, and exploration of ranked molecule choice between regression and classification models should be interesting for future *in silico* analyses. One explanation may be that underlying dataset biases (such as molecular similarity) may have consequences for classification DNNs that are different for regression models. Regardless, the data showing *Negatives Removed +SNA* rescuing model performance suggests it is reasonable to consider adding random negatives when none are available in the literature.

We also briefly explore the possibility that the method of choosing potential negative pairs assigned by SNA may have unintended consequences for ligands which are topologically similar to existing ligands. We included an option to blacklist potential molecule-target negative pairs using an alternative ligand-target prediction method, the Similarity Ensemble Approach (SEA)^8^. We expected this procedure, *SNA +SEA blacklist*, to reduce the probability of incorrectly assigning a likely ligand to be a negative example (Supplementary Information). We found that *SNA +SEA blacklist* models performed similarly to *SNA*, but leave the exploration of negative choice open for further study.

We created scrambled DNN models (*STD scrambled*; *SNA scrambled*) to serve as low-performance adversarial baselines for our experiments and evaluate them against the same hold-out benchmarks. The empirically-scrambled baseline control studies analyses yield two key observations. First, as both *STD* and *SNA* outperformed their relative scrambled controls, the DNN models here do not rely solely on memorization for their performance. Second, as *SNA* decreases the baseline down to the frequently used value for scrambled-model performance and improves performance of models on actual data, we find the *SNA* training procedure widens the predictive gap between actual and random models.

Finally, we ask, “If stochastic negatives improve model performance, how many should we impute?” Would a balanced ratio of positives-to-negatives as is common in the literature yield the highest performance, or would one more closely approximating the ligand-binding prevalence experimentally observed in comprehensive drug discovery screens produce better results? We find that for *SNA* and *Negatives Removed +SNA* models, a training dataset comprising approximately 40-60% negatives per target maximizes performance for Drug Matrix and Time Split, in support of the standard 50% negative ratio. Considering that the bulk of the improvement on Drug Matrix occurs between 10% and 40% stochastic negatives, we hypothesize that even a future Time Split with many negatives will not be drastically different from best performance ratios found with Drug Matrix.

This study is not without caveats. As noted in Methods, data from ChEMBL is biased by both researcher and assay and we have made several assumptions in aggregating datasets. We took aggregate values (median) for duplicated molecule-protein pairs to avoid over-sampling particularly well-studied pairs. We made further bulk assumptions about our dataset by asserting a single negative binding threshold (pAC50=5.0; 10 uM) when evaluating performance, agnostic to protein target. For certain proteins, a hit weaker than 10 uM may be desirable for a researcher, and for other proteins, a hit stronger than 1 nM may be the minimum affinity necessary to describe a hit. This remains an unexplored avenue of research and has interesting implications for future AUROC_r_ and AUPRC_r_ regression-as-classification analyses. Our models are additionally limited by the representation of our datasets. We did not add any structural protein information. This limits the total variance we could expect to derive from such a dataset, but we believe our method has uses where protein structural information is unavailable or where a phenotype-based readout is desired. Furthermore, our choice of the conventional ECFP4 molecular feature representation^38^ does not include information that could be obtained from 3D fingerprints or graph convolutional methods^1,39,40^.

This method is intended as an interim measure to supplement datasets while quality *in vitro* negative data may be collected and reported by experimental researchers. It is not intended as a cure-all for the lack of negative data, although it may be informative to evaluate at a finer level where experimental negatives most effectively impact model predictions, and where stochastically asserted ones are sufficient. Analysis for the particular protein target profiles that benefit under SNA conditions remain as an avenue for future study. For example, although *SNA* and *SNA +SEA blacklist* models perform similarly, highly promiscuous targets may suffer under *SNA*, and may suffer less under *SNA +SEA blacklist*. We note that SEA is a ligand-based approach and may not be a sufficiently orthogonal blacklist when considering that our neural networks are trained on ligand topology. Future studies may address this concern by incorporating biophysical models as a blacklisting methodology.

## CONCLUSION

The Stochastic Negative Addition (SNA) approach is a pharmacological data-augmentation procedure for DNNs designed to randomly assert untested negatives for public datasets where negative data are otherwise lacking. At each training epoch, new negatives are drawn to ensure that any particular negative choice does not heavily influence the model. We evaluated SNA at multiple ratios of positives-to-negatives and found that a ratio around 1:1 is optimal. We compared SNA training for both classification and regression networks trained on ChEMBL20. We found that, generally, SNA improved predictions on a held-out screening-like benchmark (Drug Matrix) with minimal effect on a 20% Time Split hold-out. Effectively, this resulted in a lower false positive rate for the screening scenario. Our random selection of negative data involved minimal computational overhead. Supplementation of DNN training with stochastic negatives provides an interim augmentation measure for datasets lacking diverse negative data until more experimental data becomes publicly available.

## METHODS

### Data Description

We filtered the ChEMBL20 database^15^ by small molecule-target affinities with a binding type “B” and reported affinity values of type IC_50_, EC_50_, K_i_, or K_d_. Adapting the ontology from Visser et al., we treat all K_i_, K_d_, IC_50_, EC_50_, and related values equivalently and broadly refer to the resulting annotations as “activity concentration 50%” (AC_50_) values^16^. We removed molecules with MW ≥ 800 Da and protein targets with fewer than 10 positive interactions. We addressed over-weighting of well-studied molecule-to-target pairs by taking the median across repeated target-molecule pairs. ChEMBL qualifies affinity using the “Relation” parameter that reports whether the true value is greater than, less than, or equal to the reported value. For all relations except “equals,” we added random noise to the values to express uncertainty (Supplementary Table 3). We transformed all AC_50_ values by –log_10_ to arrive at pAC_50_ values for training, such that pAC_50_ > 10 (i.e., < 0.1 nM) would be considered a strong binder and pAC_50_ < 5 (i.e. > 10 uM) would be considered inactive. For classification tasks, we used pAC_50_ >= 5.0 to establish positive/active class identity.

Inputs are represented as a 4096-bit RDKit Morgan Fingerprint with a radius of two^38^. Predicted values are the log transforms of affinity as described above (pAC_50_) at 2038 protein targets for each molecule.

### Data Splits

The evaluation benchmarks -- which assess two distinct use cases -- draw on Drug Matrix [CHEMBL1909046] and a 20% Time Split hold-out,^41^ and are excluded from the train and cross-validation sets. For the Time Split hold-out, we set aside approximately the final five years of ChEMBL activities, as assessed by the first reported publication date for a given interaction between molecule and protein target (see Code). Like ChEMBL, the Time Split hold-out is sparsely populated by negative data, but unlike a randomly split ChEMBL hold-out it contains more unique structures. Drug Matrix is a dataset produced by Iconix Pharmaceuticals that reports *in vitro* toxicology data for 870 chemicals across 132 protein targets^42^. Of these 132, we used the 84 targets that passed filtering steps defined in Data Description, in our training set from Drug Matrix as a way to measure how we perform on a set containing a higher ratio of negative data to positive data. Descriptions of positive and negative attributes for each split are available in Supplementary Table 4.

### Stochastic Negative Addition

Stochastic Negative Addition (*SNA*) for multi-task deep neural network training is added in an online fashion where new negative training examples for molecule-protein pairs are generated at each epoch to achieve a desired ratio of positives to negatives for each target. For the baseline *SNA* model, negatives are selected at random from all unlabeled pairs in the dataset to fulfill the desired ratio of positives to negatives at the target of interest. To evaluate the impact of potentially mis-assigning hidden positive examples during training, we developed a second method using the Similarity Ensemble Approach (SEA)^8^ to blacklist potential interactions during the sampling procedure (*SNA +SEA blacklist*). For this method, we excluded from consideration (“blacklisted”) all otherwise unlabeled pairs that achieved a positive SEA prediction with pSEA ≥ 5. We tested SNA at the following positive-to-negative ratios to find an optimal balance beginning at the baseline positive prevalence in Drug Matrix: [0.07, 0.5, 0.6, 0.75, 0.85, 0.95, 1.0, 1.33, 1.54, 2.0, 2.86, 4.0, 6.66, 10.0] .

### Negatives Removed

To assess the impact of training on purely positive data, we scrubbed all negative data (pAC_50_ < 5) from the training set (*Negatives Removed*) and evaluated it on Time Split and Drug Matrix as we did for training regimes including negatives such as *STD* and *SNA*. We applied SNA to the training regime as above to evaluate the impact of stochastic negatives on the model’s predictive ability (*Negatives Removed +SNA*). Each of the 14 ratios listed above were tested for SNA applied to models trained on the negatives-removed dataset to evaluate the impact of positive-to-negative ratios on performance.

### Software

This project was built with Python 2.7. All deep neural networks were implemented and trained in Lasagne^43^ and Theano^44^. We used RDKit for all handling of molecular structures^45^. We used NumPy^46^ and Scikit-learn^47^ for performance measures and numerical analyses, and visualizations were made with Matplotlib^48^ and Seaborn^49^.

### Multi-task deep neural network model hyperparameters and architecture

As multitask DNN performance is sensitive to architecture and hyperparameter choice, we optimized hyperparameters and architecture by considering retrospective performance on a random 20% holdout of the training dataset. We performed a grid search over varied architectures and manually explored for optimal hyperparameters. Although this optimization is not exhaustive, we focused this study on a simple representative architecture with three fully connected hidden layers with 1024, 2048 and 3072 nodes, respectively. We used an input layer of 4096 nodes, the length of our input fingerprints, and an output layer with 2038 target nodes. We use leaky rectified linear unit (leaky-ReLU) activation functions^50^ for all hidden layers and L2 weight regularization with a penalty of 5×10^−5^ and mean squared error for the loss function. We employed Stochastic Gradient Descent (SGD) with Nesterov momentum^51^ using a fixed learning rate of 0.01 and momentum of 0.4. Additionally, the hidden layers were subject to dropout^52^ with probabilities of 0.1, 0.25, 0.25 respectively.

### Model Training and Classification Accuracy Assessment

#### R-Squared

We square the correlation coefficient (r_value) from scipy.stats.linregress.

#### Area Under the Curve

Area under the curve (AUC) was analyzed for both the precision-recall curve (AUPRC) and the receiver operating characteristic curve (AUROC). For classification models, AUC was implemented as in sklearn, with a ground-truth positive threshold set to 5.0 as in training. While AUPRC and AUROC are traditionally reported for classification models, we also reported these metrics for thresholded regression models and denote these with AUROC_r_ and AUPRC_r_ for clarity. This usage has two underlying assumptions: 1) chemical screens are performed to assess hit rates past a certain biological threshold (e.g. p(AC_50_ in molar) >= 5), and higher ranking predictions from a regression model are more likely to be tested first by researchers. Given these assumptions, we post-hoc assessed regression models as classification. Prediction thresholds were chosen over the max and min of the predictions for a given model in stepsizes of 0.05 and true values were thresholded at 5.0. At each prediction threshold, true positive rate/precision, false positive rate, and recall were calculated, then the area under the curve generated from points.

### Source Code

All code necessary to reproduce this work is available at https://github.com/keiserlab/stochastic-negatives-paper under the MIT License.

## ASSOCIATED CONTENT

### Supporting Information

Supporting figures and tables include summary statistics for cross validation results and Curve Class information. The SI describes additional information important for SEA blacklisting SNA and Negatives Upweighted experiments. Figures in the SI include additional AUC, R^2^, and barplots.

## AUTHOR INFORMATION

### Author Contributions

The manuscript was written through contributions of all authors. All authors have given approval to the final version of the manuscript.

## Funding Sources

This material is based on work supported by grant number 2018-191905 from the Chan Zuckerberg Initiative DAF, an advised fund of the Silicon Valley Community Foundation (MJK) and a New Frontier Research Award from the Program for Breakthrough Biomedical Research, which is partially funded by the Sandler Foundation (MJK). ELC is supported under the National Science Foundation Graduate Research Fellowship Program under Grant No. 1650113 and is a Howard Hughes Medical Institute Gilliam Fellow.

## ACKNOWLEDGMENTS

We thank Benjamin Wong, Brian Shoichet, Garrett Gaskins, Gregory Valiant, Jason Gestwicki, Kangway Chuang, Leo Gendelev, Jessica McKinley, and Michael Mysinger for discussions and technical support.

## ABBREVIATIONS

### General Abbreviations

SNA: Stochastic Negative Addition as a procedure
AUROC: AUC of the Receiver Operating Characteristic Curve (classification)
AUPRC: AUC of the Precision-Recall Curve (classification)
AUROC_r_: AUC of the Receiver Operating Characteristic Curve (regression-as-classification)
AUPRC_r_: AUC of the Precision-Recall Curve (regression-as-classification)

### Model Abbreviations

*STD*: a “standard” model trained without SNA procedure
*STD scrambled*: *STD* model trained with y-randomization of the input training data
*SNA*: a model trained with SNA
*SNA scrambled*: *SNA* model trained with y-randomization of the input training data
*Negatives Removed*: a model trained with negatives removed from the training set
*Negatives Removed scrambled*: a *Negatives Removed* model trained with y-randomization of the input training data
*SNA +SEA blacklist*: an *SNA* model where ligands with a chance of binding (by SEA) are blacklisted from SNA choice during training.

## TOC GRAPHIC

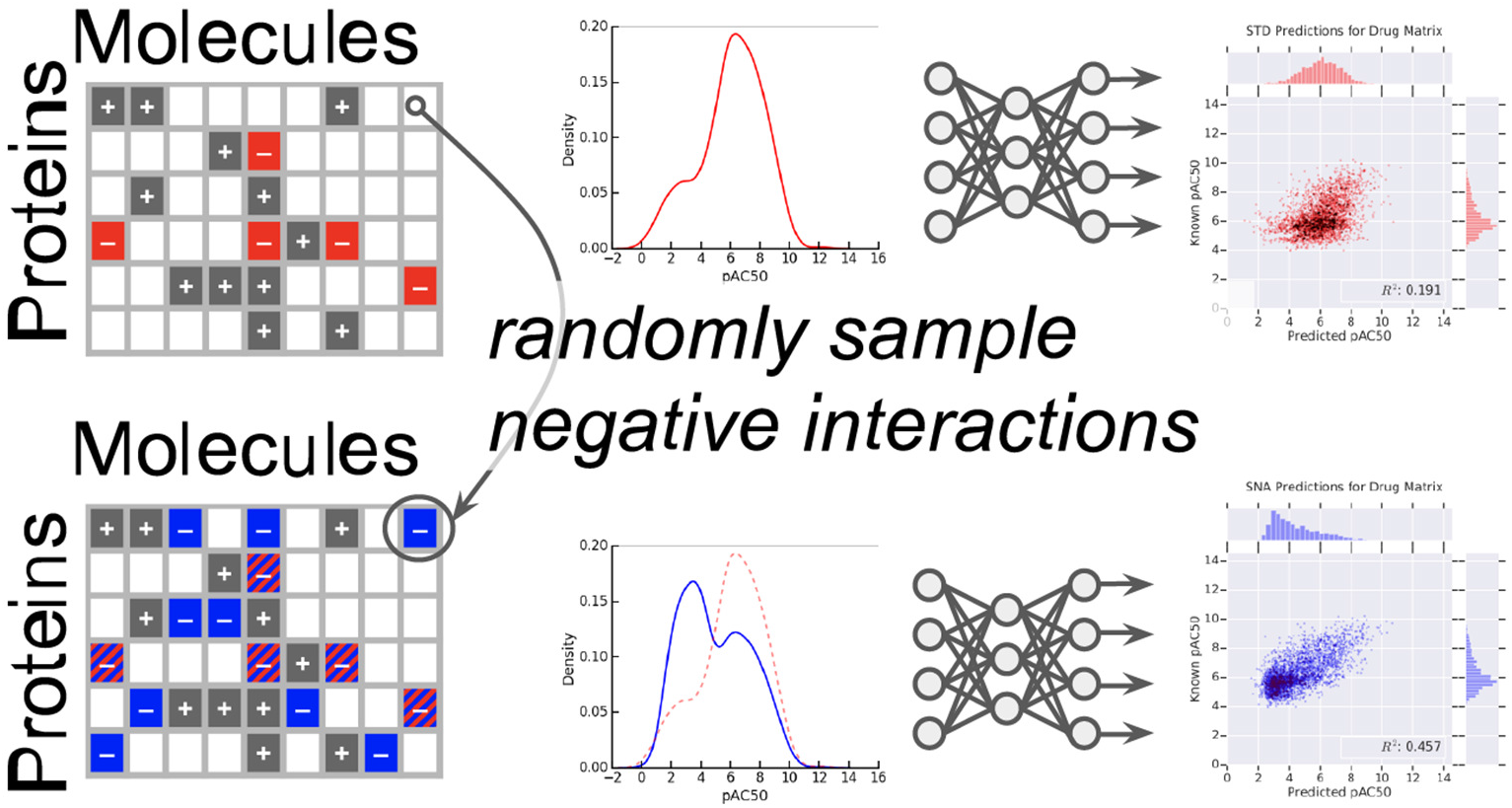

## Notes

https://github.com/keiserlab/stochastic-negatives-paper

### Competing Interest Statement

The authors have declared no competing interest.

