## Supporting Information for "Adding stochastic negative examples into machine learning improves molecular bioactivity prediction"

### Supporting Methods

#### SNA + SEA blacklisting

We were concerned over the potential impact of choosing close structurally, similar neighbors for SNA on our DNN performance. We trained an additional SNA model where predictions from Similarity Ensemble Approach (SEA) for our dataset were blacklisted if SEA predicted likely binders. Effectively, we took these predictions and did not allow the neural network to see these molecule, protein pairs as random negatives during minibatching. From the results in Supplementary Tables 1,3 and Supplementary Figures 1, 2, 5, and 6 we found that SEA blacklisting of SNA choices during training improved performance of SNA on Drug Matrix screening benchmarks and Time Split benchmarks, but the magnitude of the change was relatively small for all but the mean $R^2$ reported for Drug Matrix with regression DNNs. Classification results were also improved with very small magnitudes when SEA blacklisting was applied to SNA training. We concluded, therefore, that there exists a small risk for negative choice on overall model performance when using SNA. However, as SEA is also a ligand-based method, we suggest further research into more orthogonal methods of blacklisting.

#### Negatives Upweighted

To upweight negative examples, we calculated loss as defined in Methods then multiplied by weights based on the positive-to-negative ratio within the minibatch. We calculated a weight for known negative examples within each protein task, which is defined as the maximum value of 1.0 or the number of positive examples for the given target divided by the number of negative examples for the given target. If the weight was less than 1.0 or Nan (in the case where there are no negative examples), we set the weighting scheme to 1.0. Finally, the loss of each interaction is multiplied by the weighting scheme to upweight the impact of existing negatives.

### Supporting Figures and Tables

| Dataset | Training Type | Mean $R^2$ | $R^2$ std | Mean $AUROC_r$ | $AUROC_r$ std | Mean $AUPRC_r$ | $AUPRC_r$ std |
|---|---|---|---|---|---|---|---|
| STD | | 0.1926 | 0.0186 | 0.6886 | 0.0094 | 0.1490 | 0.0077 |
| STD scrambled | | 0.0154 | 0.0092 | 0.5538 | 0.0099 | 0.0816 | 0.0046 |
| **SNA** | | **0.4269** | **0.0272** | **0.7833** | **0.0059** | **0.4405** | **0.0079** |
| SNA scrambled | | 0.0021 | 0.0023 | 0.4842 | 0.0134 | 0.0687 | 0.0030 |
| Negatives Removed | | 0.1973 | 0.0176 | 0.6120 | 0.0076 | 0.1039 | 0.0025 |
| Negatives Removed scrambled | | 0.0065 | 0.0032 | 0.5315 | 0.0052 | 0.0756 | 0.0014 |
| **Negatives Removed +SNA** | | **0.4257** | **0.0179** | **0.7848** | **0.0053** | **0.4484** | **0.0061** |
| Negatives Upweighted | | 0.2177 | 0.0192 | 0.7024 | 0.0062 | 0.1670 | 0.0081 |
| **SNA +SEA blacklist** | Drugmatrix | **0.4411** | **0.0169** | **0.7858** | **0.0051** | **0.4528** | **0.0069** |
| **STD** | | 0.2152 | 0.0033 | **0.7388** | **0.0024** | **0.9434** | **0.0008** |
| STD scrambled | | 0.0513 | 0.0032 | 0.6340 | 0.0033 | 0.9057 | 0.0010 |
| SNA | | 0.1863 | 0.0012 | 0.7133 | 0.0025 | 0.9401 | 0.0006 |
| SNA scrambled | | 0.0020 | 0.0016 | 0.4664 | 0.0106 | 0.8540 | 0.0032 |
| **Negatives Removed** | | **0.2352** | **0.0043** | 0.7223 | 0.0050 | 0.9385 | 0.0009 |
| Negatives Removed scrambled | | 0.0547 | 0.0029 | 0.6253 | 0.0053 | 0.9024 | 0.0011 |
| Negatives Removed +SNA | | 0.1774 | 0.0018 | 0.7091 | 0.0025 | 0.9385 | 0.0007 |
| Negatives Upweighted | | 0.2179 | 0.0064 | 0.7418 | 0.0036 | 0.9444 | 0.0011 |
| SNA +SEA blacklist | Time Split | 0.1878 | 0.0021 | 0.7150 | 0.0012 | 0.9405 | 0.0004 |
| **STD** | Test | **0.6370** | **0.0041** | **0.9036** | **0.0016** | 0.9837 | 0.0004 |

| | | | | | | | |
|---|---|---|---|---|---|---|---|
| STD scrambled | | 0.0741 | 0.0026 | 0.6584 | 0.0014 | 0.9200 | 0.0019 |
| **SNA** | | **0.6428** | **0.0058** | **0.9064** | **0.0026** | **0.9848** | **0.0003** |
| SNA scrambled | | 0.0009 | 0.0004 | 0.4700 | 0.0053 | 0.8685 | 0.0012 |
| Negatives Removed | | 0.6034 | 0.0014 | 0.8362 | 0.0024 | 0.9704 | 0.0009 |
| Negatives Removed scrambled | | 0.0820 | 0.0024 | 0.6474 | 0.0011 | 0.9167 | 0.0016 |
| Negatives Removed +SNA | | 0.6026 | 0.0082 | 0.8799 | 0.0028 | 0.9795 | 0.0004 |
| **Negatives Upweighted** | | **0.6268** | **0.0057** | 0.9018 | 0.0017 | 0.9835 | 0.0003 |
| **SNA +SEA blacklist** | | **0.6462** | **0.0066** | **0.9070** | **0.0024** | **0.9849** | **0.0003** |
| **STD** | | **0.9224** | **0.0095** | **0.9809** | **0.0026** | **0.9972** | **0.0004** |
| **STD scrambled** | | **0.9212** | **0.0016** | **0.9814** | **0.0002** | **0.9973** | **0.0000** |
| SNA | | 0.8971 | 0.0100 | 0.9750 | 0.0025 | 0.9962 | 0.0004 |
| SNA scrambled | | 0.8618 | 0.0217 | 0.9725 | 0.0047 | 0.9958 | 0.0007 |
| Negatives Removed | | 0.7842 | 0.0017 | 0.8223 | 0.0012 | 0.9684 | 0.0002 |
| Negatives Removed scrambled | | 0.6454 | 0.0033 | 0.6518 | 0.0021 | 0.9283 | 0.0007 |
| Negatives Removed +SNA | | 0.7875 | 0.0129 | 0.9127 | 0.0027 | 0.9848 | 0.0005 |
| Negatives Upweighted | | 0.8712 | 0.0163 | 0.9692 | 0.0041 | 0.9954 | 0.0006 |
| SNA +SEA blacklist | Train | 0.9064 | 0.0069 | 0.9771 | 0.0015 | 0.9965 | 0.0002 |

Supplementary Table 1. Mean performance metrics and standard deviation across 5-fold cross for all regression models. Models with stochastic negatives used a 1:1 positive-to-negative ratio.

| Dataset | Training Type | Mean AUROC | AUROC std | Mean AUPRC | AUPRC std |
|---|---|---|---|---|---|
| STD (classifier) | | 0.7202 | 0.0050 | 0.1690 | 0.0024 |
| STD scrambled (classifier) | | 0.6070 | 0.0107 | 0.1075 | 0.0063 |
| **SNA (classifier)** | | **0.8168** | **0.0047** | **0.4240** | **0.0085** |
| SNA scrambled (classifier) | | 0.5645 | 0.0044 | 0.0845 | 0.0029 |
| Negatives Removed (classifier) | | 0.5434 | 0.0194 | 0.0794 | 0.0051 |
| Negatives Removed scrambled (classifier) | | 0.5275 | 0.0086 | 0.0748 | 0.0033 |
| Negatives Removed +SNA (classifier) | | 0.8035 | 0.0034 | 0.3103 | 0.0074 |
| Negatives Removed +SNA scrambled (classifier) | | 0.5440 | 0.0062 | 0.0936 | 0.0015 |
| **SNA +SEA blacklist (classifier)** | Drug Matrix | **0.8199** | **0.0034** | **0.4319** | **0.0054** |
| **STD (classifier)** | | **0.7314** | **0.0044** | **0.9397** | **0.0010** |
| STD scrambled (classifier) | | 0.6955 | 0.0088 | 0.9259 | 0.0028 |
| SNA (classifier) | | 0.7010 | 0.0016 | 0.9346 | 0.0006 |
| SNA scrambled (classifier) | | 0.6579 | 0.0029 | 0.9144 | 0.0022 |
| Negatives Removed (classifier) | | 0.6332 | 0.0100 | 0.9099 | 0.0037 |
| Negatives Removed scrambled (classifier) | | 0.6262 | 0.0056 | 0.9069 | 0.0024 |
| Negatives Removed +SNA (classifier) | | 0.6542 | 0.0031 | 0.9187 | 0.0011 |
| Negatives Removed +SNA scrambled (classifier) | | 0.5739 | 0.0055 | 0.8893 | 0.0018 |
| SNA +SEA blacklist (classifier) | Time Split | 0.7031 | 0.0018 | 0.9354 | 0.0005 |
| **STD (classifier)** | | **0.9044** | **0.0019** | **0.9827** | **0.0001** |
| STD scrambled (classifier) | | 0.7401 | 0.0038 | 0.9419 | 0.0022 |
| **SNA (classifier)** | | 0.9010 | 0.0014 | **0.9823** | **0.0003** |
| SNA scrambled (classifier) | Test | 0.7354 | 0.0035 | 0.9407 | 0.0012 |

| | | | | | |
|---|---|---|---|---|---|
| Negatives Removed (classifier) | | 0.6642 | 0.0075 | 0.9238 | 0.0026 |
| Negatives Removed scrambled (classifier) | | 0.6255 | 0.0088 | 0.9111 | 0.0014 |
| Negatives Removed +SNA (classifier) | | 0.7091 | 0.0048 | 0.9343 | 0.0020 |
| Negatives Removed +SNA scrambled (classifier) | | 0.6233 | 0.0065 | 0.9110 | 0.0014 |
| SNA +SEA blacklist (classifier) | | 0.9004 | 0.0011 | 0.9822 | 0.0003 |
| **STD (classifier)** | | **0.9606** | **0.0033** | **0.9937** | **0.0006** |
| STD scrambled (classifier) | | 0.8162 | 0.0110 | 0.9665 | 0.0028 |
| **SNA (classifier)** | | **0.9652** | **0.0023** | **0.9944** | **0.0004** |
| SNA scrambled (classifier) | | 0.7473 | 0.0021 | 0.9433 | 0.0016 |
| Negatives Removed (classifier) | | 0.6655 | 0.0106 | 0.9241 | 0.0026 |
| Negatives Removed scrambled (classifier) | | 0.6264 | 0.0052 | 0.9113 | 0.0019 |
| Negatives Removed +SNA (classifier) | | 0.7178 | 0.0028 | 0.9359 | 0.0008 |
| Negatives Removed +SNA scrambled (classifier) | | 0.6232 | 0.0047 | 0.9110 | 0.0017 |
| SNA +SEA blacklist (classifier) | Train | 0.9593 | 0.0010 | 0.9933 | 0.0002 |

Supplementary Table 2. Mean performance metrics and standard deviation across 5-fold cross for all classification models. Models with stochastic negatives used a 1:1 positive-to-negative ratio.

| ChEMBL activity | Dataset action action |
| --- | --- |
| =' or '<' | accept pIC50 value as-is |
| >' | add np.random 2-3 logs to reported pIC50 |

Supplementary Table 3: ChEMBL activity relation actions.

| Dataset | Num Positives | Num Negatives | Total | Percent Positive | Percent Negative |
|---|---|---|---|---|---|
| Train All | 403409 | 154826 | 558235 | 72.26508549 | 27.73491451 |
| Test Fold 0 | 80861 | 31341 | 112202 | 72.06734283 | 27.93265717 |
| Train Fold 0 | 322548 | 123485 | 446033 | 72.31482872 | 27.68517128 |
| Test Fold 1 | 81373 | 32890 | 114263 | 71.21552909 | 28.78447091 |
| Train Fold 1 | 322036 | 121936 | 443972 | 72.53520492 | 27.46479508 |
| Test Fold 2 | 79566 | 29624 | 109190 | 72.86931038 | 27.13068962 |
| Train Fold 2 | 323843 | 125202 | 449045 | 72.11816188 | 27.88183812 |
| Test Fold 3 | 80312 | 29874 | 110186 | 72.88766268 | 27.11233732 |
| Train Fold 3 | 323097 | 124952 | 448049 | 72.11197882 | 27.88802118 |
| Test Fold 4 | 81297 | 31097 | 112394 | 72.33215296 | 27.66784704 |
| Train Fold 4 | 322112 | 123729 | 445841 | 72.24817816 | 27.75182184 |
| Time Split | 83155 | 33774 | 116929 | 71.11580532 | 28.88419468 |
| Drug Matrix (dose response) | 2714 | 330 | 3044 | 89.15900131 | 10.84099869 |
| Drug Matrix (primary) | 2714 | 35451 | 38165 | 7.111227565 | 92.88877244 |

Supplementary Table 4: Positive and Negative splits for Test, Train, Time Split, and Drug Matrix

| Dataset | Model | Positive to Negative Ratio | Target Minimum Negative Percent | Target Percent Positive | AUROC | AUROC std | AUPRC | AUPRC std | R2 | R2 std |
|---|---|---|---|---|---|---|---|---|---|---|
| | | 0.0000 | 0.0000 | 100.0000 | 0.6120 | 0.0076 | 0.1039 | 0.0025 | 0.1973 | 0.0176 |
| | | 0.0753 | 92.9973 | 7.0027 | 0.7404 | 0.0059 | 0.3937 | 0.0043 | 0.3529 | 0.0139 |
| | | 0.1111 | 90.0009 | 9.9991 | 0.7495 | 0.0053 | 0.4061 | 0.0073 | 0.3654 | 0.0207 |
| | | 0.2500 | 80.0000 | 20.0000 | 0.7671 | 0.0060 | 0.4270 | 0.0070 | 0.3865 | 0.0171 |
| | | 0.4286 | 69.9986 | 30.0014 | 0.7708 | 0.0061 | 0.4308 | 0.0085 | 0.3912 | 0.0193 |
| | | 0.6666 | **60.0024** | **39.9976** | 0.7757 | 0.0104 | **0.4403** | **0.0104** | **0.4060** | **0.0185** |
| | | 0.8182 | **54.9995** | **45.0005** | **0.7864** | **0.0082** | **0.4456** | **0.0087** | **0.4245** | **0.0249** |
| | | 1.0000 | **50.0000** | **50.0000** | **0.7848** | **0.0053** | **0.4479** | **0.0062** | **0.4257** | **0.0179** |
| | | 1.2222 | **45.0005** | **54.9995** | **0.7937** | **0.0078** | **0.4534** | **0.0099** | **0.4327** | **0.0226** |
| | | 1.5000 | **40.0000** | **60.0000** | **0.7969** | **0.0115** | **0.4528** | **0.0106** | **0.4317** | **0.0202** |
| | | 2.3333 | **30.0003** | **69.9997** | 0.7798 | 0.0053 | 0.3695 | 0.0186 | **0.4321** | **0.0139** |
| | | 4.0000 | 20.0000 | 80.0000 | 0.7358 | 0.0055 | 0.2457 | 0.0165 | 0.3966 | 0.0163 |
| | | 9.0000 | 10.0000 | 90.0000 | 0.6482 | 0.0070 | 0.1195 | 0.0052 | 0.2816 | 0.0150 |
| | Negatives Removed +SNA | 19.0000 | 5.0000 | 95.0000 | 0.6124 | 0.0077 | 0.1042 | 0.0054 | 0.2033 | 0.0087 |
| | | 0.0000 | 0.0000 | 100.0000 | 0.6886 | 0.0094 | 0.1490 | 0.0077 | 0.1926 | 0.0186 |
| | | 0.0753 | 92.9973 | 7.0027 | 0.7474 | 0.0103 | 0.4032 | 0.0070 | 0.3641 | 0.0215 |
| | | 0.1111 | 90.0009 | 9.9991 | 0.7540 | 0.0098 | 0.4166 | 0.0087 | 0.3800 | 0.0202 |
| Drug Matrix | SNA | 0.2500 | 80.0000 | 20.0000 | 0.7697 | 0.0070 | 0.4380 | 0.0048 | 0.4028 | 0.0202 |

| | | | | | | | | | | |
|---|---|---|---|---|---|---|---|---|---|---|
| | | **0.4286** | **69.9986** | **30.0014** | 0.7721 | 0.0078 | 0.4367 | 0.0057 | **0.4092** | **0.0203** |
| | | **0.6666** | **60.0024** | **39.9976** | 0.7799 | 0.0034 | **0.4433** | **0.0018** | **0.4187** | **0.0166** |
| | | **0.8182** | **54.9995** | **45.0005** | **0.7857** | **0.0064** | **0.4453** | **0.0040** | **0.4239** | **0.0166** |
| | | **1.0000** | **50.0000** | **50.0000** | **0.7841** | **0.0061** | **0.4428** | **0.0054** | **0.4282** | **0.0215** |
| | | **1.2222** | **45.0005** | **54.9995** | **0.7901** | **0.0032** | **0.4502** | **0.0024** | **0.4329** | **0.0151** |
| | | **1.5000** | **40.0000** | **60.0000** | **0.7908** | **0.0032** | **0.4490** | **0.0062** | **0.4341** | **0.0159** |
| | | **2.3333** | **30.0003** | **69.9997** | 0.7803 | 0.0066 | 0.3702 | 0.0194 | **0.4383** | **0.0194** |
| | | 4.0000 | 20.0000 | 80.0000 | 0.7434 | 0.0066 | 0.2504 | 0.0159 | 0.3987 | 0.0093 |
| | | 9.0000 | 10.0000 | 90.0000 | 0.7012 | 0.0064 | 0.1661 | 0.0065 | 0.3184 | 0.0152 |
| | | 19.0000 | 5.0000 | 95.0000 | 0.6831 | 0.0063 | 0.1477 | 0.0042 | 0.2172 | 0.0229 |
| | | 0.0000 | 0.0000 | 100.0000 | 0.8362 | 0.0024 | 0.9704 | 0.0009 | 0.6034 | 0.0014 |
| | | 0.0753 | 92.9973 | 7.0027 | 0.8511 | 0.0021 | 0.9741 | 0.0003 | 0.4798 | 0.0053 |
| | | 0.1111 | 90.0009 | 9.9991 | 0.8600 | 0.0023 | 0.9760 | 0.0003 | 0.5137 | 0.0066 |
| | | 0.2500 | 80.0000 | 20.0000 | 0.8730 | 0.0025 | 0.9784 | 0.0004 | 0.5631 | 0.0072 |
| | | **0.4286** | **69.9986** | **30.0014** | **0.8782** | **0.0024** | **0.9793** | **0.0002** | 0.5852 | 0.0055 |
| | | **0.6666** | **60.0024** | **39.9976** | **0.8790** | **0.0027** | **0.9795** | **0.0004** | 0.5910 | 0.0085 |
| | | **0.8182** | **54.9995** | **45.0005** | **0.8779** | **0.0027** | **0.9792** | **0.0004** | 0.5966 | 0.0051 |
| | | **1.0000** | **50.0000** | **50.0000** | **0.8799** | **0.0028** | **0.9795** | **0.0004** | 0.6026 | 0.0082 |
| Test | Negatives Removed +SNA | **1.2222** | **45.0005** | **54.9995** | **0.8801** | **0.0033** | **0.9796** | **0.0005** | 0.6036 | 0.0079 |

| | | | | | | | | | |
|---|---|---|---|---|---|---|---|---|---|
| | **1.5000** | **40.0000** | **60.0000** | **0.8793** | **0.0018** | **0.9794** | **0.0004** | 0.6040 | 0.0053 |
| | **2.3333** | **30.0003** | **69.9997** | 0.8745 | **0.0035** | **0.9783** | **0.0010** | **0.6203** | **0.0032** |
| | **4.0000** | **20.0000** | **80.0000** | 0.8649 | 0.0024 | 0.9764 | 0.0007 | **0.6205** | **0.0019** |
| | 9.0000 | 10.0000 | 90.0000 | 0.8465 | 0.0023 | 0.9727 | 0.0008 | 0.6084 | 0.0021 |
| | 19.0000 | 5.0000 | 95.0000 | 0.8358 | 0.0028 | 0.9705 | 0.0008 | 0.6017 | 0.0024 |
| | 0.0000 | 0.0000 | 100.0000 | 0.9036 | 0.0016 | 0.9837 | 0.0004 | 0.6370 | 0.0041 |
| | 0.0753 | 92.9973 | 7.0027 | 0.8674 | 0.0020 | 0.9778 | 0.0002 | 0.5122 | 0.0044 |
| | 0.1111 | 90.0009 | 9.9991 | 0.8799 | 0.0024 | 0.9802 | 0.0002 | 0.5521 | 0.0063 |
| | 0.2500 | 80.0000 | 20.0000 | 0.8962 | 0.0022 | 0.9831 | 0.0002 | 0.6074 | 0.0068 |
| | 0.4286 | 69.9986 | 30.0014 | 0.9022 | 0.0028 | 0.9841 | 0.0003 | 0.6277 | 0.0068 |
| | 0.6666 | 60.0024 | 39.9976 | 0.9035 | 0.0027 | 0.9843 | 0.0002 | 0.6328 | 0.0066 |
| | **0.8182** | **54.9995** | **45.0005** | **0.9055** | **0.0024** | **0.9846** | **0.0003** | 0.6408 | 0.0049 |
| | **1.0000** | **50.0000** | **50.0000** | **0.9061** | **0.0025** | **0.9847** | **0.0003** | **0.6419** | **0.0054** |
| | **1.2222** | **45.0005** | **54.9995** | **0.9064** | **0.0023** | **0.9848** | **0.0003** | **0.6432** | **0.0066** |
| | **1.5000** | **40.0000** | **60.0000** | **0.9070** | **0.0027** | **0.9849** | **0.0002** | **0.6451** | **0.0076** |
| | **2.3333** | **30.0003** | **69.9997** | **0.9085** | **0.0012** | **0.9849** | **0.0003** | **0.6520** | **0.0034** |
| | **4.0000** | **20.0000** | **80.0000** | **0.9076** | **0.0019** | **0.9846** | **0.0004** | **0.6516** | **0.0039** |
| | **9.0000** | **10.0000** | **90.0000** | **0.9069** | **0.0027** | **0.9844** | **0.0004** | **0.6509** | **0.0051** |
| SNA | **19.0000** | **5.0000** | **95.0000** | **0.9066** | **0.0024** | 0.9842 | 0.0005 | **0.6501** | **0.0043** |

| | | | | | | | | | | |
|---|---|---|---|---|---|---|---|---|---|---|
| | | **0.0000** | **0.0000** | **100.0000** | **0.7223** | **0.0050** | **0.9385** | **0.0009** | **0.2352** | **0.0043** |
| | | 0.0753 | 92.9973 | 7.0027 | 0.7025 | 0.0016 | 0.9360 | 0.0004 | 0.1608 | 0.0013 |
| | | 0.1111 | 90.0009 | 9.9991 | 0.7011 | 0.0008 | 0.9359 | 0.0002 | 0.1616 | 0.0013 |
| | | 0.2500 | 80.0000 | 20.0000 | 0.7032 | 0.0019 | 0.9367 | 0.0004 | 0.1677 | 0.0011 |
| | | 0.4286 | 69.9986 | 30.0014 | 0.7028 | 0.0011 | 0.9367 | 0.0002 | 0.1691 | 0.0017 |
| | | 0.6666 | 60.0024 | 39.9976 | 0.7049 | 0.0014 | 0.9374 | 0.0004 | 0.1720 | 0.0016 |
| | | 0.8182 | 54.9995 | 45.0005 | 0.7070 | 0.0023 | 0.9379 | 0.0006 | 0.1764 | 0.0018 |
| | | 1.0000 | 50.0000 | 50.0000 | 0.7091 | 0.0025 | 0.9385 | 0.0007 | 0.1774 | 0.0018 |
| | | 1.2222 | 45.0005 | 54.9995 | 0.7098 | 0.0026 | 0.9387 | 0.0006 | 0.1793 | 0.0035 |
| | | 1.5000 | 40.0000 | 60.0000 | 0.7111 | 0.0019 | 0.9390 | 0.0004 | 0.1792 | 0.0012 |
| | | 2.3333 | 30.0003 | 69.9997 | 0.6973 | 0.0012 | 0.9356 | 0.0004 | 0.1820 | 0.0027 |
| | | 4.0000 | 20.0000 | 80.0000 | 0.7005 | 0.0025 | 0.9347 | 0.0007 | 0.1899 | 0.0052 |
| | | 9.0000 | 10.0000 | 90.0000 | 0.7166 | 0.0021 | 0.9380 | 0.0007 | 0.2163 | 0.0040 |
| | Negatives Removed +SNA | **19.0000** | **5.0000** | **95.0000** | **0.7277** | **0.0056** | **0.9398** | **0.0012** | **0.2437** | **0.0061** |
| | | **0.0000** | **0.0000** | **100.0000** | **0.7388** | **0.0024** | **0.9434** | **0.0008** | **0.2152** | **0.0033** |
| | | 0.0753 | 92.9973 | 7.0027 | 0.7015 | 0.0012 | 0.9369 | 0.0004 | 0.1682 | 0.0011 |
| | | 0.1111 | 90.0009 | 9.9991 | 0.7037 | 0.0015 | 0.9377 | 0.0004 | 0.1731 | 0.0015 |
| | | 0.2500 | 80.0000 | 20.0000 | 0.7060 | 0.0020 | 0.9383 | 0.0006 | 0.1785 | 0.0017 |
| Time Split | SNA | 0.4286 | 69.9986 | 30.0014 | 0.7090 | 0.0016 | 0.9391 | 0.0005 | 0.1825 | 0.0028 |

| | | | | | | | | | | |
|---|---|---|---|---|---|---|---|---|---|---|
| | | 0.6666 | 60.0024 | 39.9976 | 0.7104 | 0.0033 | 0.9392 | 0.0007 | 0.1835 | 0.0018 |
| | | 0.8182 | 54.9995 | 45.0005 | 0.7113 | 0.0023 | 0.9395 | 0.0006 | 0.1846 | 0.0024 |
| | | 1.0000 | 50.0000 | 50.0000 | 0.7109 | 0.0023 | 0.9395 | 0.0006 | 0.1831 | 0.0028 |
| | | 1.2222 | 45.0005 | 54.9995 | 0.7122 | 0.0029 | 0.9398 | 0.0007 | 0.1838 | 0.0022 |
| | | 1.5000 | 40.0000 | 60.0000 | 0.7129 | 0.0023 | 0.9400 | 0.0006 | 0.1843 | 0.0023 |
| | | 2.3333 | 30.0003 | 69.9997 | 0.7131 | 0.0033 | 0.9398 | 0.0007 | 0.1911 | 0.0024 |
| | | 4.0000 | 20.0000 | 80.0000 | 0.7210 | 0.0022 | 0.9406 | 0.0005 | 0.1997 | 0.0036 |
| | | **9.0000** | **10.0000** | **90.0000** | **0.7329** | **0.0018** | **0.9426** | **0.0004** | **0.2154** | **0.0021** |
| | | **19.0000** | **5.0000** | **95.0000** | **0.7380** | **0.0041** | **0.9431** | **0.0011** | **0.2213** | **0.0050** |
| | | 0.0000 | 0.0000 | 100.0000 | 0.8223 | 0.0012 | 0.9684 | 0.0002 | 0.7842 | 0.0017 |
| | | 0.0753 | 92.9973 | 7.0027 | 0.8775 | 0.0014 | 0.9787 | 0.0002 | 0.5620 | 0.0036 |
| | | 0.1111 | 90.0009 | 9.9991 | 0.8905 | 0.0015 | 0.9811 | 0.0003 | 0.6236 | 0.0043 |
| | | **0.2500** | **80.0000** | **20.0000** | **0.9090** | **0.0017** | 0.9843 | 0.0003 | 0.7247 | 0.0037 |
| | | **0.4286** | **69.9986** | **30.0014** | **0.9154** | **0.0017** | **0.9853** | **0.0003** | 0.7744 | 0.0097 |
| | | **0.6666** | **60.0024** | **39.9976** | **0.9158** | **0.0028** | **0.9854** | **0.0004** | **0.7835** | **0.0118** |
| | | **0.8182** | **54.9995** | **45.0005** | **0.9095** | **0.0020** | 0.9842 | 0.0004 | 0.7726 | 0.0108 |
| | | **1.0000** | **50.0000** | **50.0000** | **0.9127** | **0.0027** | **0.9848** | **0.0005** | **0.7875** | **0.0129** |
| | | **1.2222** | **45.0005** | **54.9995** | **0.9123** | **0.0038** | **0.9847** | **0.0006** | **0.7887** | **0.0166** |
| Train | Negatives Removed +SNA | **1.5000** | **40.0000** | **60.0000** | **0.9104** | **0.0017** | 0.9843 | 0.0003 | 0.7840 | 0.0088 |

| | | | | | | | | |
|---|---|---|---|---|---|---|---|---|
| | **2.3333** | **30.0003** | **69.9997** | 0.8950 | 0.0036 | 0.9817 | 0.0005 | **0.8049** | **0.0133** |
| | **4.0000** | **20.0000** | **80.0000** | 0.8756 | 0.0016 | 0.9783 | 0.0003 | **0.8074** | **0.0062** |
| | **9.0000** | **10.0000** | **90.0000** | 0.8450 | 0.0007 | 0.9727 | 0.0002 | **0.7918** | **0.0085** |
| | **19.0000** | **5.0000** | **95.0000** | 0.8240 | 0.0021 | 0.9688 | 0.0005 | **0.7840** | **0.0039** |
| | **0.0000** | **0.0000** | **100.0000** | **0.9809** | **0.0026** | 0.9972 | 0.0004 | **0.9224** | **0.0095** |
| | 0.0753 | 92.9973 | 7.0027 | 0.8966 | 0.0006 | 0.9830 | 0.0001 | 0.5984 | 0.0011 |
| | 0.1111 | 90.0009 | 9.9991 | 0.9156 | 0.0007 | 0.9864 | 0.0002 | 0.6670 | 0.0021 |
| | 0.2500 | 80.0000 | 20.0000 | 0.9462 | 0.0009 | 0.9916 | 0.0002 | 0.7859 | 0.0040 |
| | 0.4286 | 69.9986 | 30.0014 | 0.9623 | 0.0010 | 0.9942 | 0.0001 | 0.8487 | 0.0028 |
| | 0.6666 | 60.0024 | 39.9976 | 0.9673 | 0.0009 | 0.9950 | 0.0001 | 0.8687 | 0.0025 |
| | 0.8182 | 54.9995 | 45.0005 | 0.9733 | 0.0012 | 0.9959 | 0.0002 | 0.8912 | 0.0048 |
| | 1.0000 | 50.0000 | 50.0000 | 0.9753 | 0.0017 | 0.9962 | 0.0003 | 0.8984 | 0.0061 |
| | 1.2222 | 45.0005 | 54.9995 | 0.9746 | 0.0018 | 0.9961 | 0.0003 | 0.8961 | 0.0074 |
| | 1.5000 | 40.0000 | 60.0000 | 0.9762 | 0.0016 | 0.9964 | 0.0002 | 0.9022 | 0.0075 |
| | **2.3333** | **30.0003** | **69.9997** | **0.9832** | **0.0017** | 0.9975 | 0.0003 | **0.9287** | **0.0062** |
| | **4.0000** | **20.0000** | **80.0000** | **0.9861** | **0.0007** | **0.9979** | **0.0001** | **0.9399** | **0.0032** |
| | **9.0000** | **10.0000** | **90.0000** | **0.9844** | **0.0026** | **0.9977** | **0.0004** | **0.9339** | **0.0097** |
| SNA | **19.0000** | **5.0000** | **95.0000** | **0.9830** | **0.0042** | **0.9974** | **0.0006** | **0.9296** | **0.0165** |

Supplementary Table 5: Regression performance for SNA models across multiple positive to negative ratios.

| Dataset | Model | Positive to Negative Ratio | Target Minimum Negative Percent | Target Percent Positive | AUROC | AUROC std | AUPRC | AUPRC std |
|---|---|---|---|---|---|---|---|---|
| | Negatives Removed +SNA (classifier | 0.0000 | 0.0000 | 100.0000 | 0.5434 | 0.0194 | 0.0794 | 0.0051 |
| | | 0.0753 | 92.9973 | 7.0027 | 0.7391 | 0.0022 | 0.2235 | 0.0012 |
| | | 0.1111 | 90.0009 | 9.9991 | 0.7515 | 0.0020 | 0.2570 | 0.0016 |
| | | 0.2500 | 80.0000 | 20.0000 | 0.7787 | 0.0089 | 0.2667 | 0.0158 |
| | | 0.4286 | 69.9986 | 30.0014 | 0.7934 | 0.0026 | 0.2940 | 0.0066 |
| | | **0.6666** | **60.0024** | **39.9976** | 0.8003 | 0.0025 | **0.3035** | **0.0038** |
| | | **0.8182** | **54.9995** | **45.0005** | 0.8014 | 0.0034 | **0.3138** | **0.0048** |
| | | **1.0000** | **50.0000** | **50.0000** | **0.8035** | **0.0034** | **0.3103** | **0.0074** |
| | | **1.2222** | **45.0005** | **54.9995** | **0.8064** | **0.0037** | **0.3116** | **0.0065** |
| | | **1.5000** | **40.0000** | **60.0000** | 0.8086 | 0.0035 | **0.3060** | **0.0091** |
| | | 2.3333 | 30.0003 | 69.9997 | 0.7690 | 0.0040 | 0.2832 | 0.0104 |
| | | 4.0000 | 20.0000 | 80.0000 | 0.6846 | 0.0041 | 0.1558 | 0.0120 |
| | | 9.0000 | 10.0000 | 90.0000 | 0.5331 | 0.0188 | 0.0834 | 0.0060 |
| | | 19.0000 | 5.0000 | 95.0000 | 0.5466 | 0.0059 | 0.0777 | 0.0012 |
| Drug Matrix | SNA (classifier) | 0.0000 | 0.0000 | 100.0000 | 0.7202 | 0.0050 | 0.1690 | 0.0024 |
| | | 0.0753 | 92.9973 | 7.0027 | 0.7870 | 0.0034 | 0.3258 | 0.0038 |
| | | 0.1111 | 90.0009 | 9.9991 | 0.7952 | 0.0048 | 0.3485 | 0.0051 |
| | | 0.2500 | 80.0000 | 20.0000 | 0.8095 | 0.0028 | 0.4038 | 0.0059 |
| | | 0.4286 | 69.9986 | 30.0014 | 0.8113 | 0.0038 | 0.4159 | 0.0050 |
| | | **0.6666** | **60.0024** | **39.9976** | 0.8167 | 0.0017 | **0.4235** | **0.0041** |
| | | **0.8182** | **54.9995** | **45.0005** | 0.8119 | 0.0040 | **0.4176** | **0.0085** |
| | | **1.0000** | **50.0000** | **50.0000** | **0.8168** | **0.0047** | **0.4240** | **0.0085** |
| | | **1.2222** | **45.0005** | **54.9995** | **0.8216** | **0.0042** | **0.4280** | **0.0064** |

| | | | | | | | | |
|---|---|---|---|---|---|---|---|---|
| | | **1.5000** | **40.0000** | **60.0000** | **0.8228** | **0.0038** | **0.4248** | **0.0041** |
| | | 2.3333 | 30.0003 | 69.9997 | 0.8051 | 0.0025 | 0.3924 | 0.0074 |
| | | 4.0000 | 20.0000 | 80.0000 | 0.7789 | 0.0033 | 0.3115 | 0.0105 |
| | | 9.0000 | 10.0000 | 90.0000 | 0.7470 | 0.0028 | 0.2151 | 0.0104 |
| | | 19.0000 | 5.0000 | 95.0000 | 0.7136 | 0.0019 | 0.1637 | 0.0041 |
| | Negatives Removed +SNA (classifier | 0.0000 | 0.0000 | 100.0000 | 0.6642 | 0.0075 | 0.9238 | 0.0026 |
| | | 0.0753 | 92.9973 | 7.0027 | 0.6342 | 0.0021 | 0.9226 | 0.0020 |
| | | 0.1111 | 90.0009 | 9.9991 | 0.6713 | 0.0017 | 0.9273 | 0.0016 |
| | | 0.2500 | 80.0000 | 20.0000 | 0.6930 | 0.0044 | 0.9306 | 0.0019 |
| | | **0.4286** | **69.9986** | **30.0014** | 0.7018 | 0.0033 | **0.9324** | **0.0010** |
| | | **0.6666** | **60.0024** | **39.9976** | **0.7057** | **0.0033** | **0.9332** | **0.0022** |
| | | **0.8182** | **54.9995** | **45.0005** | **0.7085** | **0.0028** | **0.9340** | **0.0014** |
| | | **1.0000** | **50.0000** | **50.0000** | **0.7091** | **0.0048** | **0.9343** | **0.0020** |
| | | **1.2222** | **45.0005** | **54.9995** | **0.7097** | **0.0034** | **0.9344** | **0.0015** |
| | | **1.5000** | **40.0000** | **60.0000** | **0.7073** | **0.0040** | **0.9338** | **0.0024** |
| | | 2.3333 | 30.0003 | 69.9997 | 0.6774 | 0.0045 | 0.9296 | 0.0016 |
| | | 4.0000 | 20.0000 | 80.0000 | 0.6177 | 0.0059 | 0.9122 | 0.0028 |
| | | 9.0000 | 10.0000 | 90.0000 | 0.5692 | 0.0154 | 0.8976 | 0.0034 |
| | | 19.0000 | 5.0000 | 95.0000 | 0.6538 | 0.0073 | 0.9201 | 0.0016 |
| test | SNA (classifier) | **0.0000** | **0.0000** | **100.0000** | **0.9044** | **0.0019** | 0.9827 | 0.0001 |
| | | 0.0753 | 92.9973 | 7.0027 | 0.8074 | 0.0017 | 0.9626 | 0.0005 |
| | | 0.1111 | 90.0009 | 9.9991 | 0.8297 | 0.0066 | 0.9677 | 0.0014 |
| | | 0.2500 | 80.0000 | 20.0000 | 0.8706 | 0.0019 | 0.9763 | 0.0004 |
| | | 0.4286 | 69.9986 | 30.0014 | 0.8881 | 0.0019 | 0.9799 | 0.0003 |
| | | 0.6666 | 60.0024 | 39.9976 | 0.8926 | 0.0018 | 0.9807 | 0.0002 |

| | | | | | | | | |
|---|---|---|---|---|---|---|---|---|
| | | 0.8182 | 54.9995 | 45.0005 | 0.8999 | 0.0015 | 0.9821 | 0.0003 |
| | | 1.0000 | 50.0000 | 50.0000 | 0.9010 | 0.0014 | 0.9823 | 0.0003 |
| | | **1.2222** | **45.0005** | **54.9995** | **0.8993** | **0.0072** | **0.9820** | **0.0014** |
| | | **1.5000** | **40.0000** | **60.0000** | **0.9030** | **0.0013** | **0.9827** | **0.0004** |
| | | **2.3333** | **30.0003** | **69.9997** | **0.9063** | **0.0013** | **0.9833** | **0.0004** |
| | | **4.0000** | **20.0000** | **80.0000** | **0.9057** | **0.0024** | **0.9831** | **0.0004** |
| | | **9.0000** | **10.0000** | **90.0000** | **0.9052** | **0.0023** | **0.9829** | **0.0003** |
| | | **19.0000** | **5.0000** | **95.0000** | **0.9031** | **0.0032** | 0.9825 | 0.0003 |
| | | 0.0000 | 0.0000 | 100.0000 | 0.6332 | 0.0100 | 0.9099 | 0.0037 |
| | | 0.0753 | 92.9973 | 7.0027 | 0.5904 | 0.0007 | 0.9036 | 0.0006 |
| | | 0.1111 | 90.0009 | 9.9991 | 0.6339 | 0.0009 | 0.9134 | 0.0003 |
| | | 0.2500 | 80.0000 | 20.0000 | 0.6454 | 0.0037 | 0.9168 | 0.0015 |
| | | **0.4286** | **69.9986** | **30.0014** | 0.6534 | 0.0018 | **0.9192** | **0.0003** |
| | | **0.6666** | **60.0024** | **39.9976** | **0.6554** | **0.0019** | **0.9193** | **0.0007** |
| | | **0.8182** | **54.9995** | **45.0005** | **0.6569** | **0.0013** | **0.9197** | **0.0003** |
| | | **1.0000** | **50.0000** | **50.0000** | **0.6542** | **0.0031** | **0.9187** | **0.0011** |
| | | **1.2222** | **45.0005** | **54.9995** | **0.6574** | **0.0013** | **0.9194** | **0.0001** |
| | | **1.5000** | **40.0000** | **60.0000** | **0.6546** | **0.0028** | **0.9189** | **0.0006** |
| | | 2.3333 | 30.0003 | 69.9997 | 0.6247 | 0.0015 | 0.9123 | 0.0008 |
| | | 4.0000 | 20.0000 | 80.0000 | 0.5893 | 0.0054 | 0.8951 | 0.0017 |
| | | 9.0000 | 10.0000 | 90.0000 | 0.5745 | 0.0064 | 0.8916 | 0.0041 |
| | Negatives Removed +SNA (classifier) | 19.0000 | 5.0000 | 95.0000 | 0.6259 | 0.0086 | 0.9058 | 0.0035 |
| | | **0.0000** | **0.0000** | **100.0000** | **0.7314** | **0.0044** | **0.9397** | **0.0010** |
| | | 0.0753 | 92.9973 | 7.0027 | 0.6791 | 0.0012 | 0.9281 | 0.0003 |
| timesplit | SNA (classifier) | 0.1111 | 90.0009 | 9.9991 | 0.6829 | 0.0010 | 0.9294 | 0.0003 |

| | | | | | | | | |
|---|---|---|---|---|---|---|---|---|
| | | 0.2500 | 80.0000 | 20.0000 | 0.6890 | 0.0006 | 0.9320 | 0.0003 |
| | | 0.4286 | 69.9986 | 30.0014 | 0.6954 | 0.0007 | 0.9336 | 0.0002 |
| | | 0.6666 | 60.0024 | 39.9976 | 0.6950 | 0.0011 | 0.9333 | 0.0005 |
| | | 0.8182 | 54.9995 | 45.0005 | 0.6982 | 0.0026 | 0.9338 | 0.0007 |
| | | 1.0000 | 50.0000 | 50.0000 | 0.7010 | 0.0016 | 0.9346 | 0.0006 |
| | | 1.2222 | 45.0005 | 54.9995 | 0.7029 | 0.0040 | 0.9350 | 0.0012 |
| | | 1.5000 | 40.0000 | 60.0000 | 0.7012 | 0.0023 | 0.9343 | 0.0008 |
| | | 2.3333 | 30.0003 | 69.9997 | 0.7134 | 0.0037 | 0.9374 | 0.0012 |
| | | 4.0000 | 20.0000 | 80.0000 | 0.7226 | 0.0026 | 0.9387 | 0.0009 |
| | | **9.0000** | **10.0000** | **90.0000** | **0.7302** | **0.0031** | **0.9398** | **0.0007** |
| | | **19.0000** | **5.0000** | **95.0000** | **0.7321** | **0.0022** | **0.9401** | **0.0006** |
| | | 0.0000 | 0.0000 | 100.0000 | 0.6655 | 0.0106 | 0.9241 | 0.0026 |
| | | 0.0753 | 92.9973 | 7.0027 | 0.6363 | 0.0019 | 0.9232 | 0.0008 |
| | | 0.1111 | 90.0009 | 9.9991 | 0.6750 | 0.0009 | 0.9281 | 0.0007 |
| | | 0.2500 | 80.0000 | 20.0000 | 0.7001 | 0.0022 | 0.9321 | 0.0009 |
| | | 0.4286 | 69.9986 | 30.0014 | 0.7087 | 0.0040 | 0.9337 | 0.0013 |
| | | **0.6666** | **60.0024** | **39.9976** | 0.7138 | 0.0027 | **0.9349** | **0.0007** |
| | | **0.8182** | **54.9995** | **45.0005** | **0.7169** | **0.0005** | **0.9356** | **0.0005** |
| | | **1.0000** | **50.0000** | **50.0000** | **0.7178** | **0.0028** | **0.9359** | **0.0008** |
| | | **1.2222** | **45.0005** | **54.9995** | **0.7184** | **0.0017** | **0.9361** | **0.0008** |
| | | **1.5000** | **40.0000** | **60.0000** | **0.7153** | **0.0039** | **0.9352** | **0.0009** |
| | | 2.3333 | 30.0003 | 69.9997 | 0.6831 | 0.0039 | 0.9308 | 0.0014 |
| | | 4.0000 | 20.0000 | 80.0000 | 0.6204 | 0.0043 | 0.9126 | 0.0015 |
| | | 9.0000 | 10.0000 | 90.0000 | 0.5712 | 0.0151 | 0.8981 | 0.0037 |
| train | Negatives Removed +SNA (classifier) | 19.0000 | 5.0000 | 95.0000 | 0.6552 | 0.0086 | 0.9204 | 0.0026 |

| | | | | | | | | |
|---|---|---|---|---|---|---|---|---|
| | | **0.0000** | **0.0000** | **100.0000** | **0.9606** | **0.0033** | **0.9937** | **0.0006** |
| | | 0.0753 | 92.9973 | 7.0027 | 0.8277 | 0.0009 | 0.9669 | 0.0002 |
| | | 0.1111 | 90.0009 | 9.9991 | 0.8547 | 0.0075 | 0.9728 | 0.0017 |
| | | 0.2500 | 80.0000 | 20.0000 | 0.9101 | 0.0011 | 0.9841 | 0.0002 |
| | | 0.4286 | 69.9986 | 30.0014 | 0.9387 | 0.0009 | 0.9896 | 0.0002 |
| | | **0.6666** | **60.0024** | **39.9976** | **0.9471** | **0.0012** | **0.9912** | **0.0002** |
| | | **0.8182** | **54.9995** | **45.0005** | **0.9640** | **0.0009** | **0.9942** | **0.0002** |
| | | **1.0000** | **50.0000** | **50.0000** | **0.9652** | **0.0023** | **0.9944** | **0.0004** |
| | | **1.2222** | **45.0005** | **54.9995** | **0.9586** | **0.0176** | **0.9931** | **0.0033** |
| | | **1.5000** | **40.0000** | **60.0000** | **0.9698** | **0.0020** | **0.9952** | **0.0003** |
| | | **2.3333** | **30.0003** | **69.9997** | **0.9674** | **0.0032** | **0.9948** | **0.0005** |
| | | **4.0000** | **20.0000** | **80.0000** | **0.9648** | **0.0027** | **0.9944** | **0.0005** |
| | | **9.0000** | **10.0000** | **90.0000** | **0.9619** | **0.0016** | **0.9939** | **0.0003** |
| | SNA (classifier) | **19.0000** | **5.0000** | **95.0000** | **0.9596** | **0.0054** | **0.9935** | **0.0010** |

Supplementary Table 6: Classification performance for SNA models across multiple positive to negative ratios.

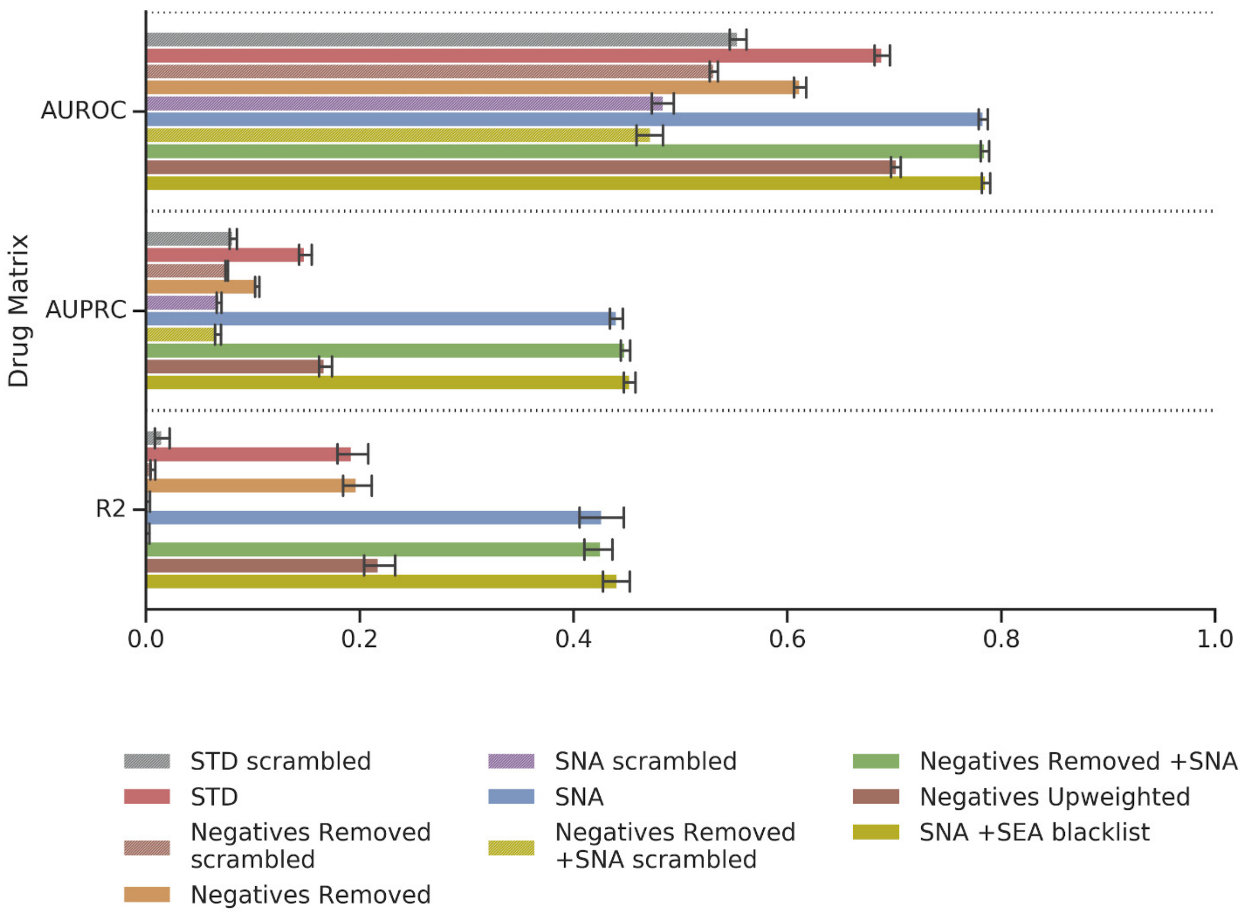

Supplementary Figure 1. Drug Matrix performance for all regression models. Note: R2 values for *SNA scrambled* and *Negatives Removed +SNA* are close to 0.0

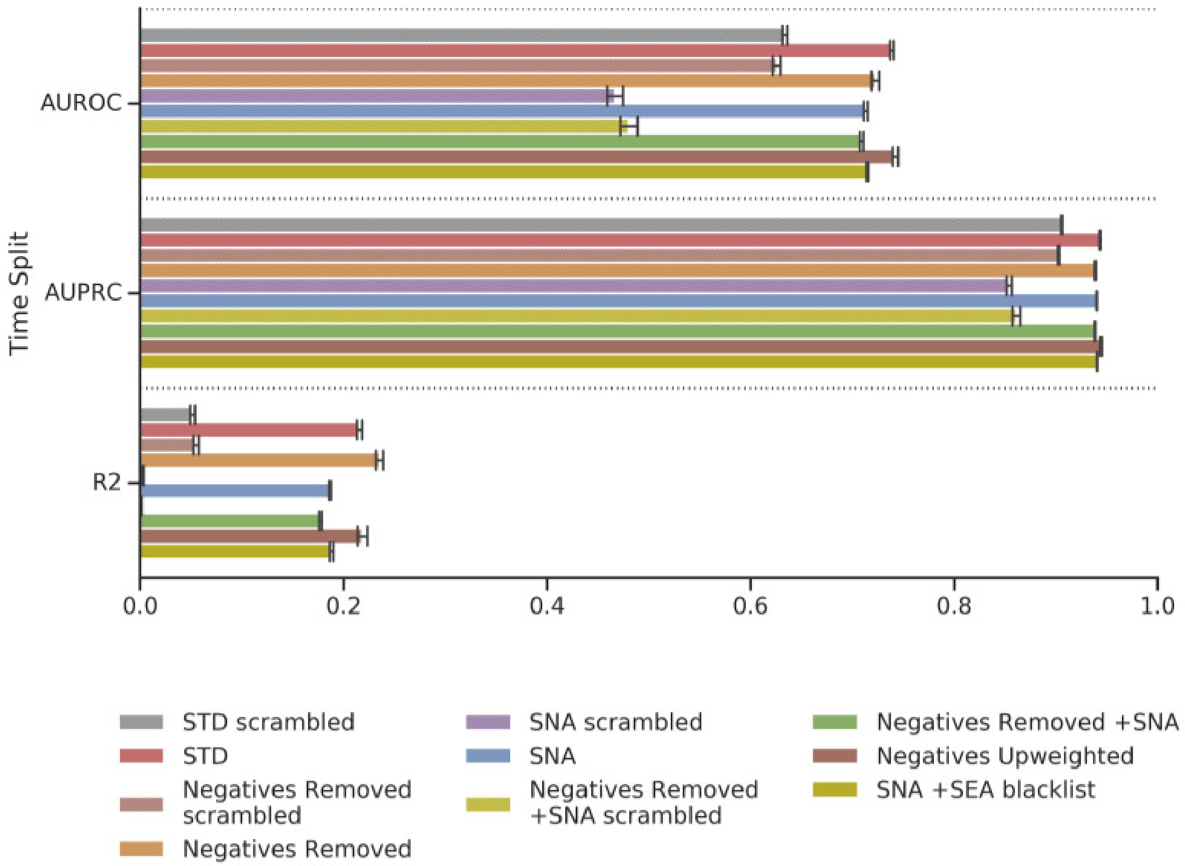

Supplementary Figure 2. Time Split performance for all regression models. Note: R2 values for *SNA scrambled* and *Negatives Removed +SNA* are close to 0.0

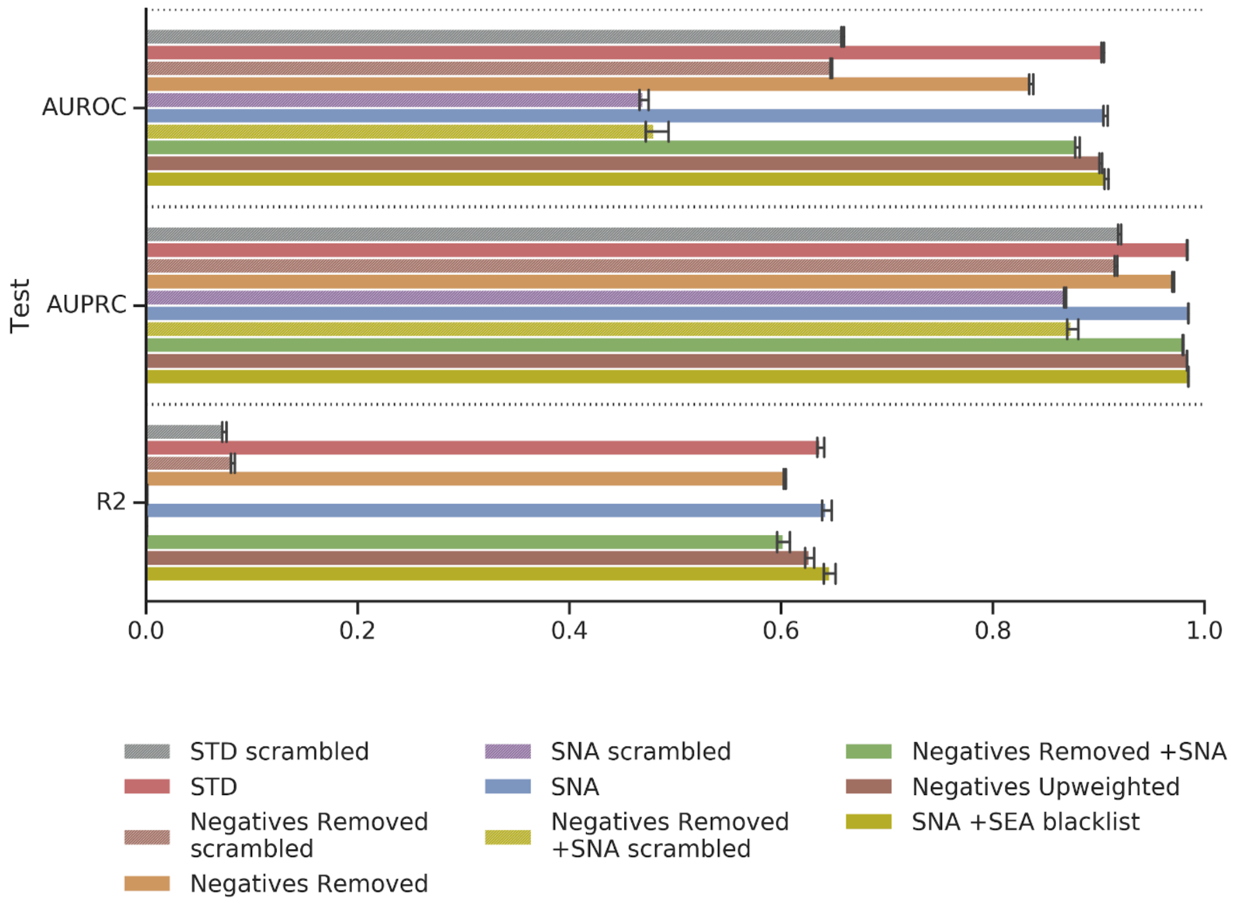

Supplementary Figure 3. Train performance for all regression models.

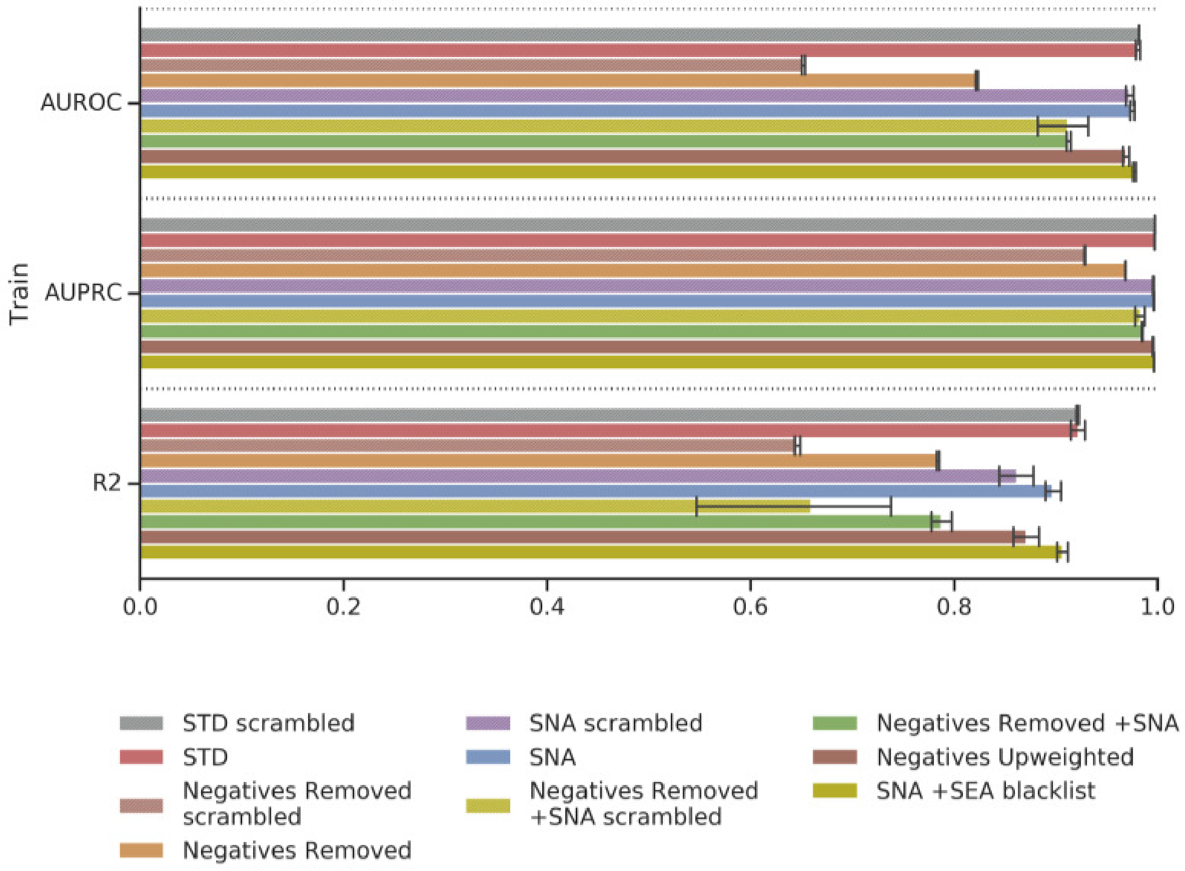

Supplementary Figure 4. Test performance for all regression models.

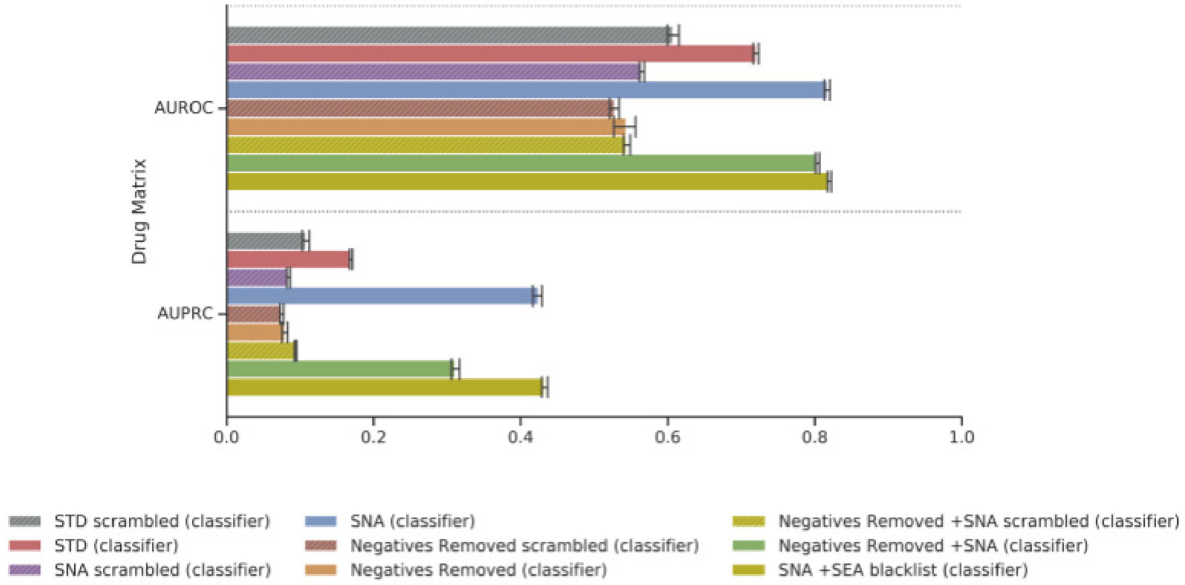

Supplementary Figure 5. Drug Matrix performance for all classification models.

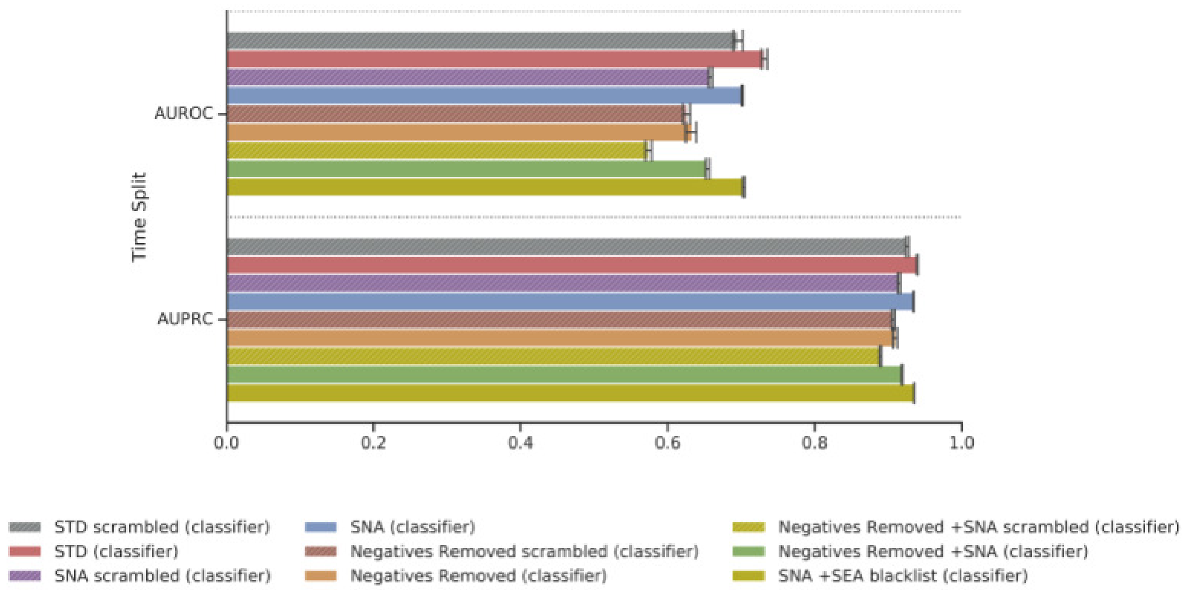

Supplementary Figure 6. Time Split performance for all classification models.

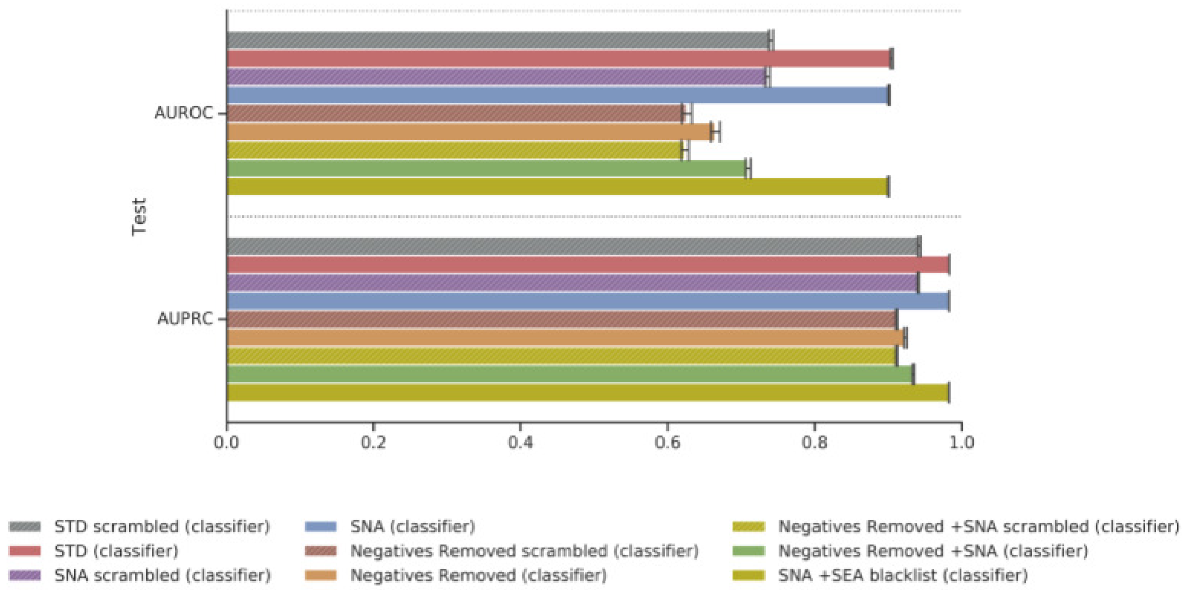

Supplementary Figure 7. Test performance for all classification models.

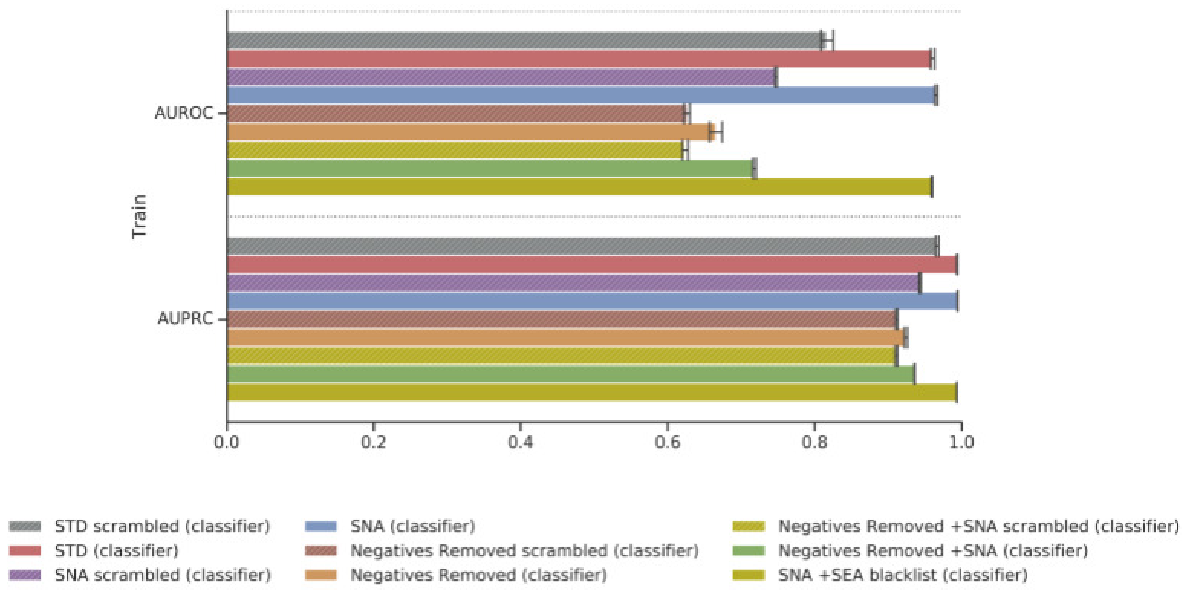

Supplementary Figure 8. Train performance for all classification models.

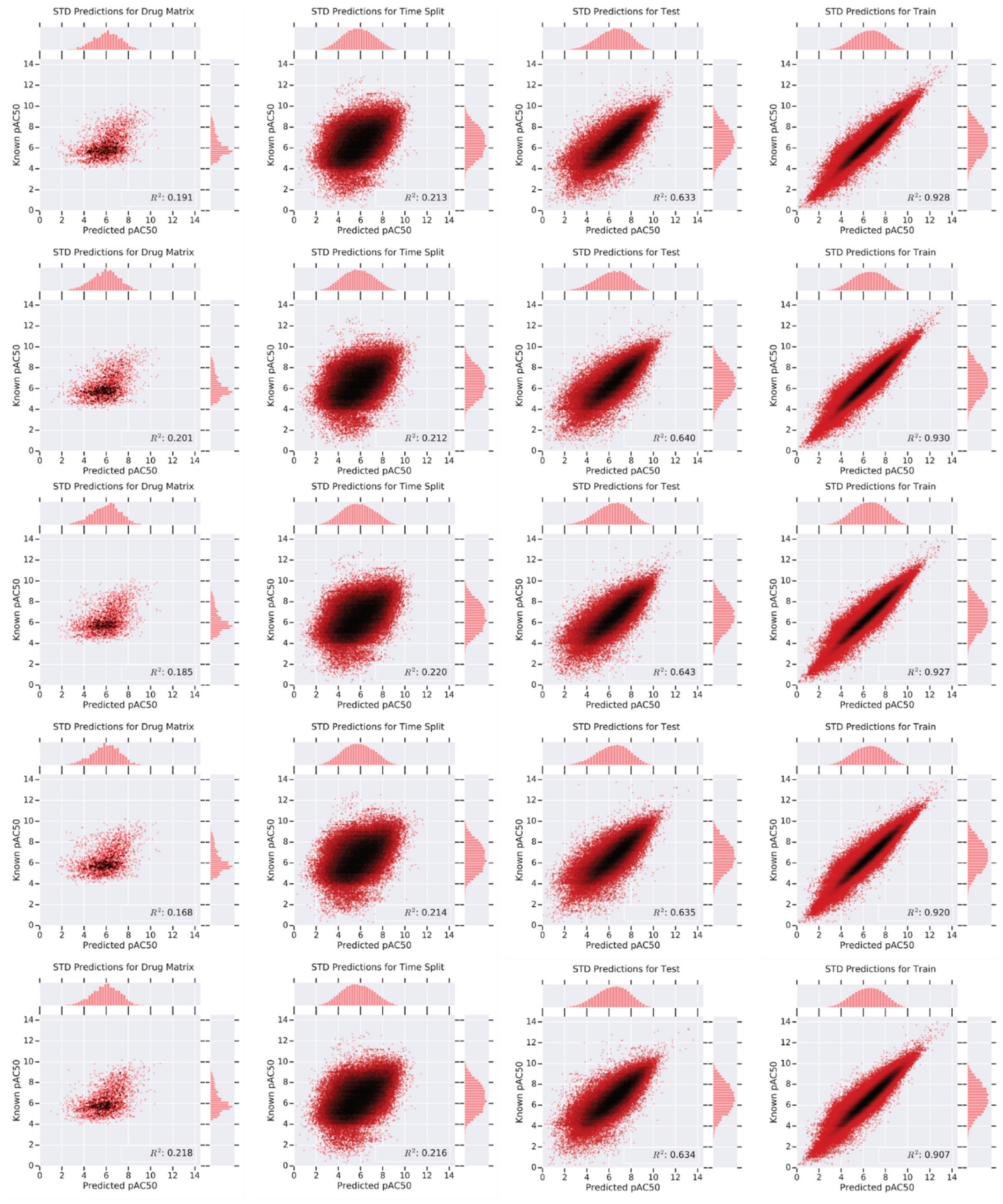

Supplementary Figure 9. *STD* model $R^2$ plots for Drug Matrix (column 1), Time Split (column 2), Test (column 3), and Train (column 4) across each fold (0-4, top to bottom, increasing).

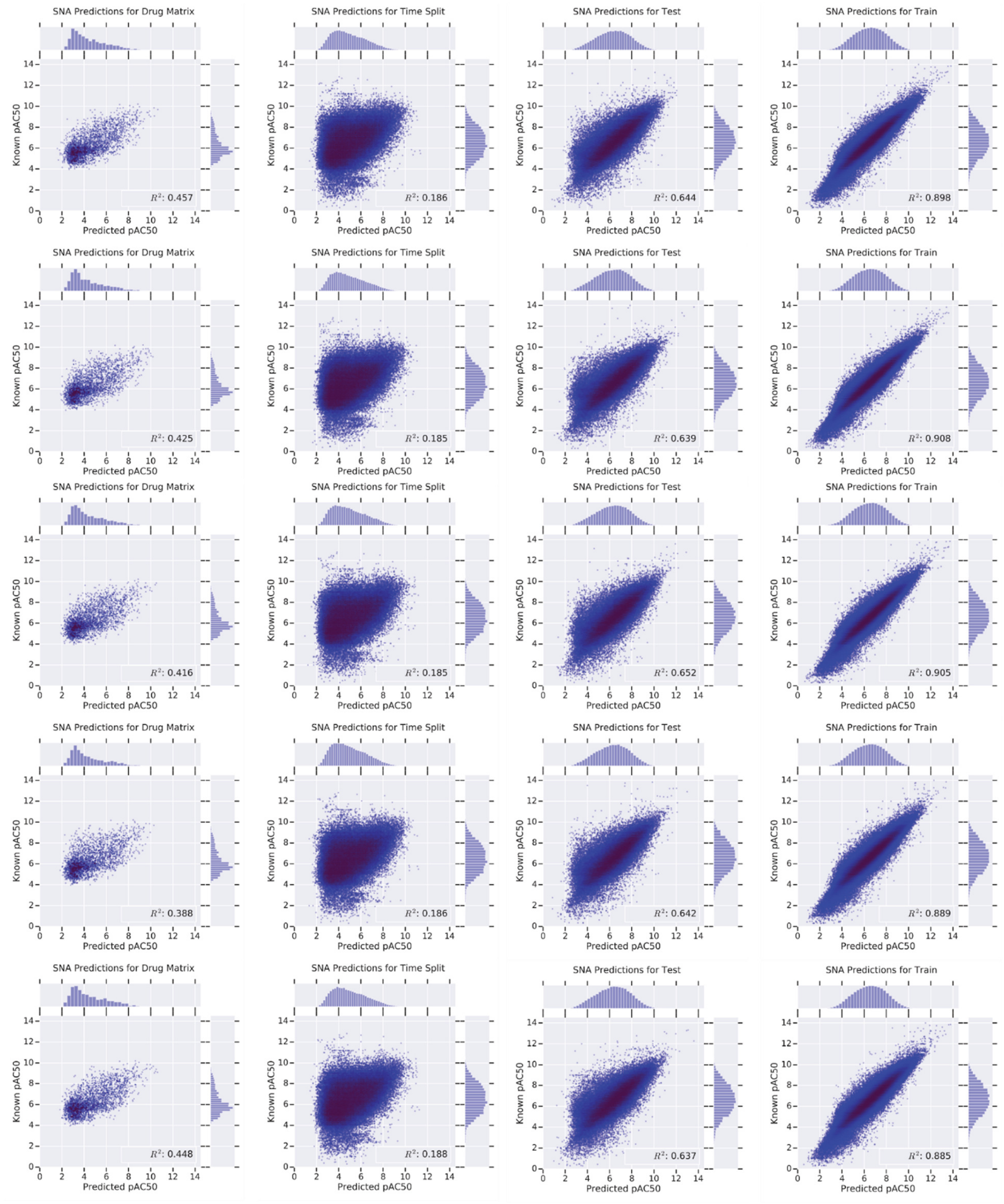

Supplementary Figure 10. *SNA* DNN $R^2$ plots for Drug Matrix (column 1), Time Split (column 2), Test (column 3), and Train (column 4) across each fold (0-4, top to bottom, increasing).

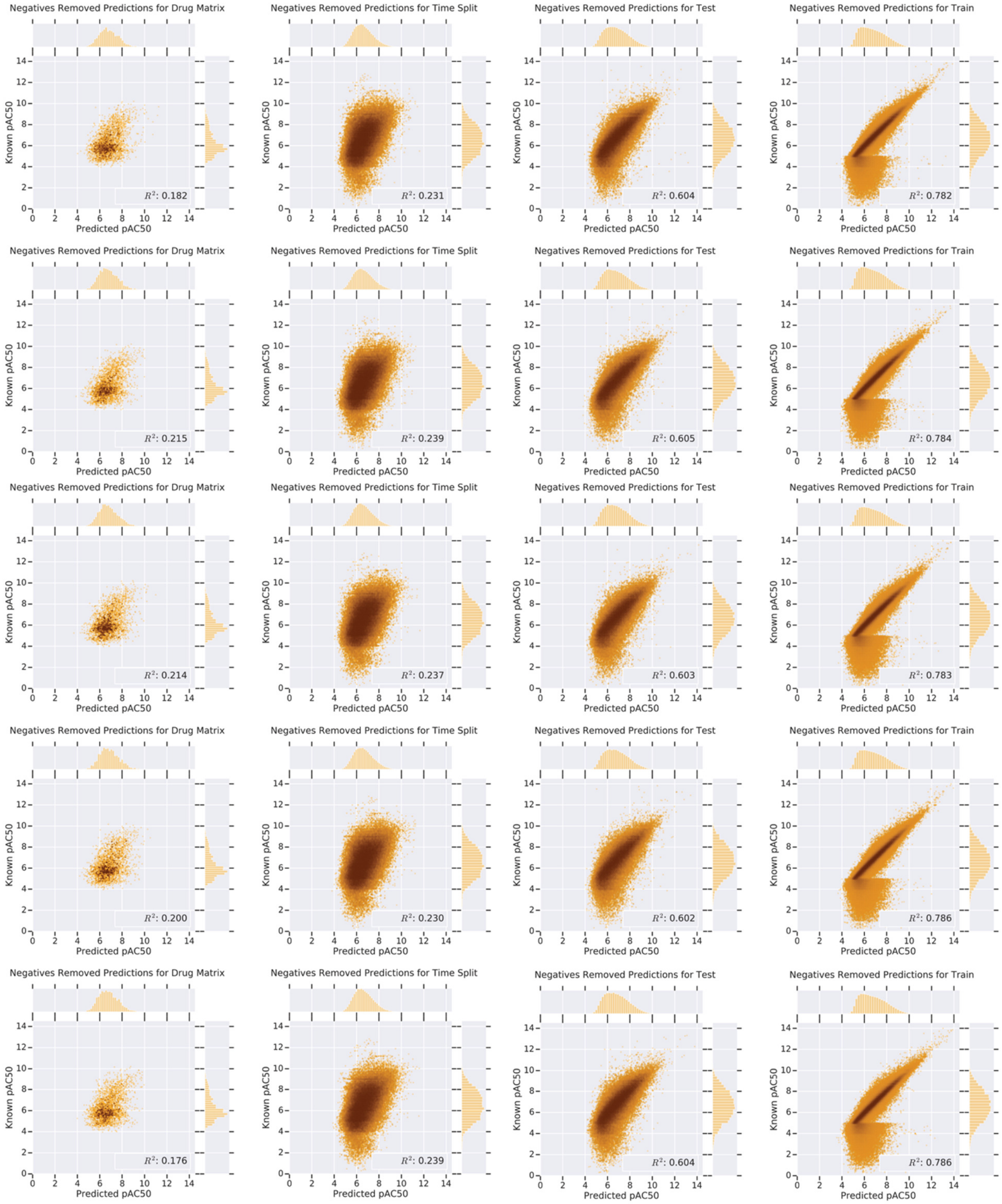

Supplementary Figure 11. *Negatives Removed* DNN $R^2$ plots for Drug Matrix (column 1), Time Split (column 2), Test (column 3), and Train (column 4) across each fold (0-4, top to bottom, increasing).

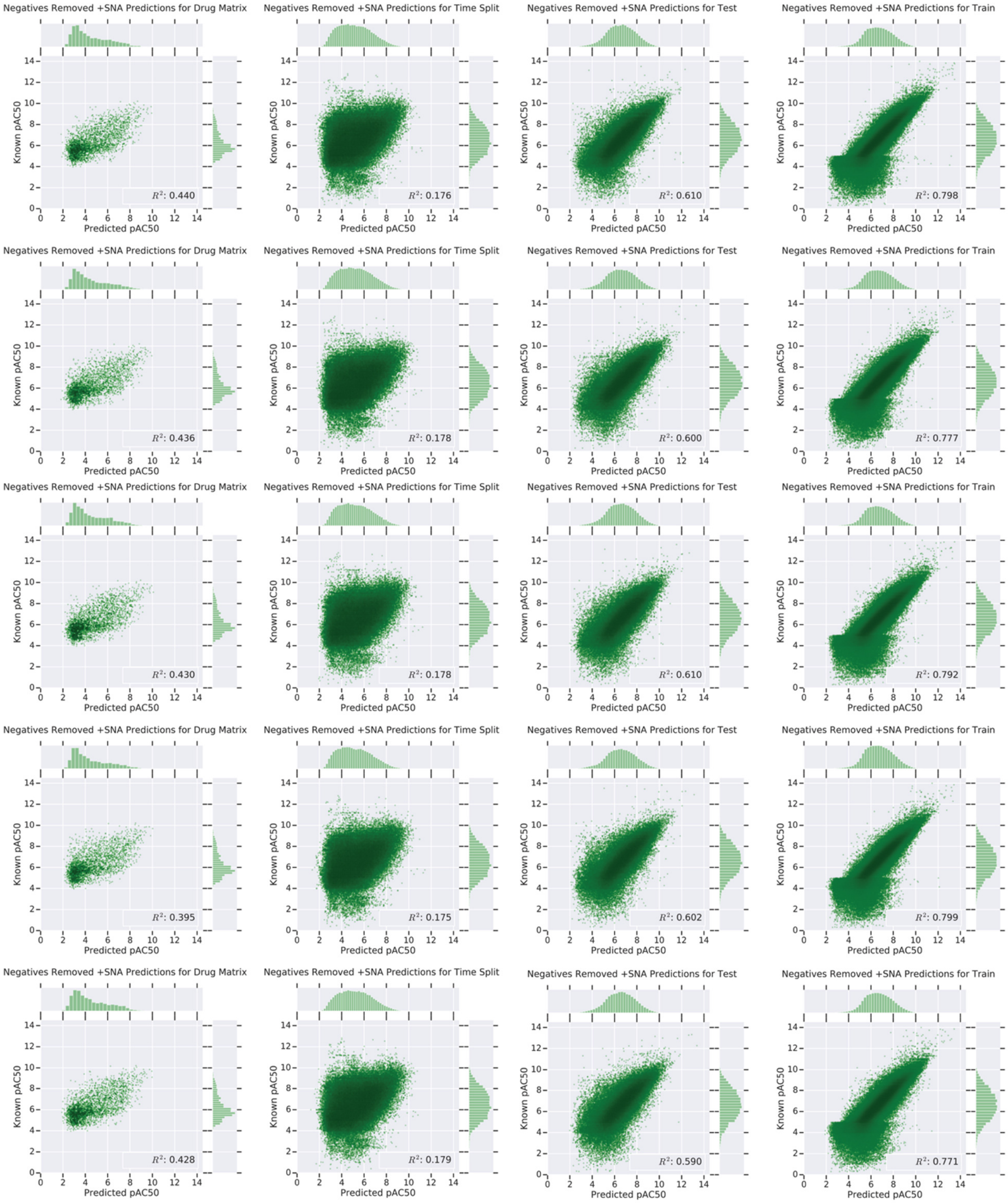

Supplementary Figure 12. *Negatives Removed +SNA* DNN $R^2$ plots for Drug Matrix (column 1), Time Split (column 2), Test (column 3), and Train (column 4) across each fold (0-4, top to bottom, increasing).

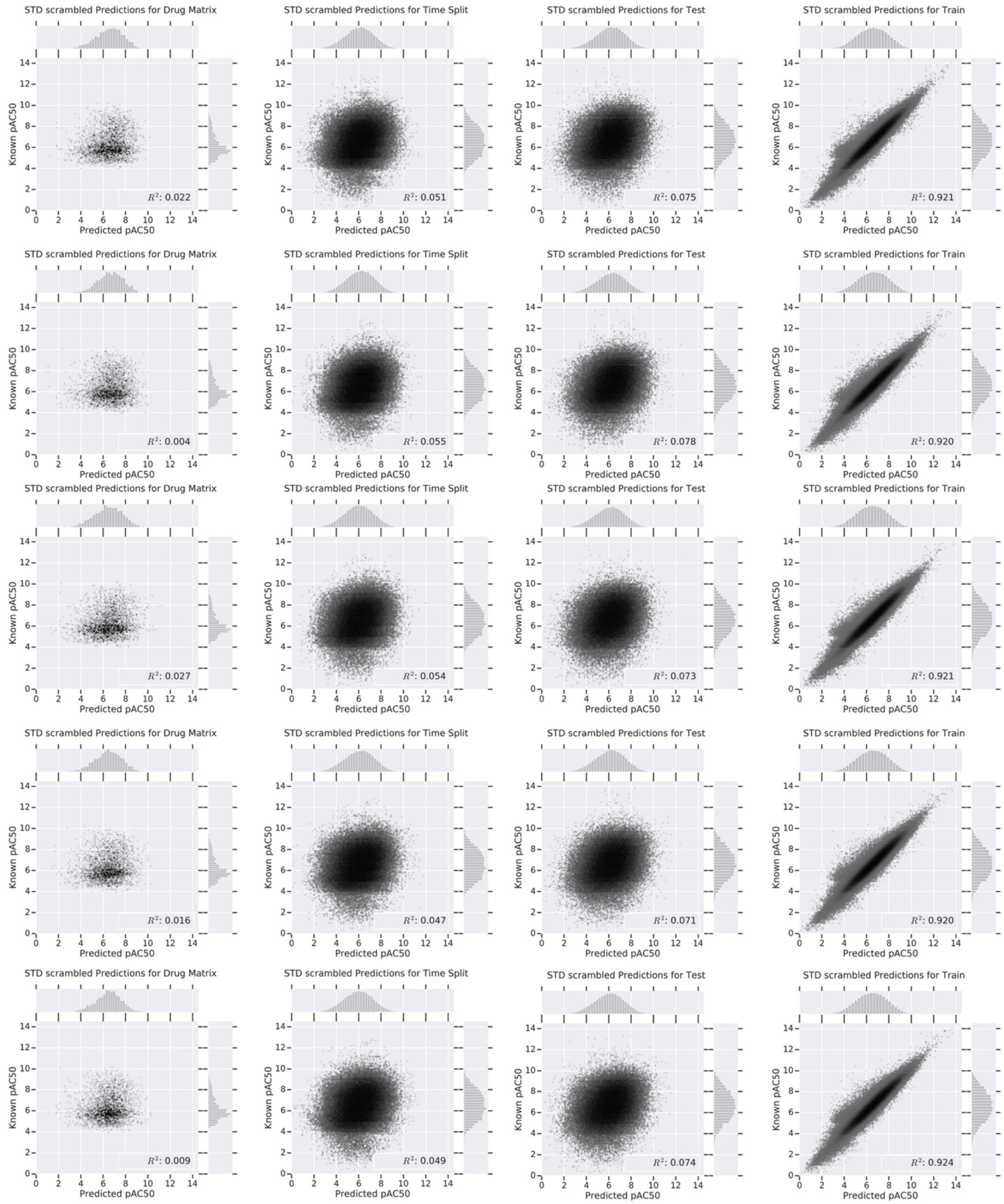

Supplementary Figure 13. *STD scrambled* (y-randomized training set control) DNN $R^2$ plots for Drug Matrix (column 1), Time Split (column 2), Test (column 3), and Train (column 4) across each fold (0-4, top to bottom, increasing).

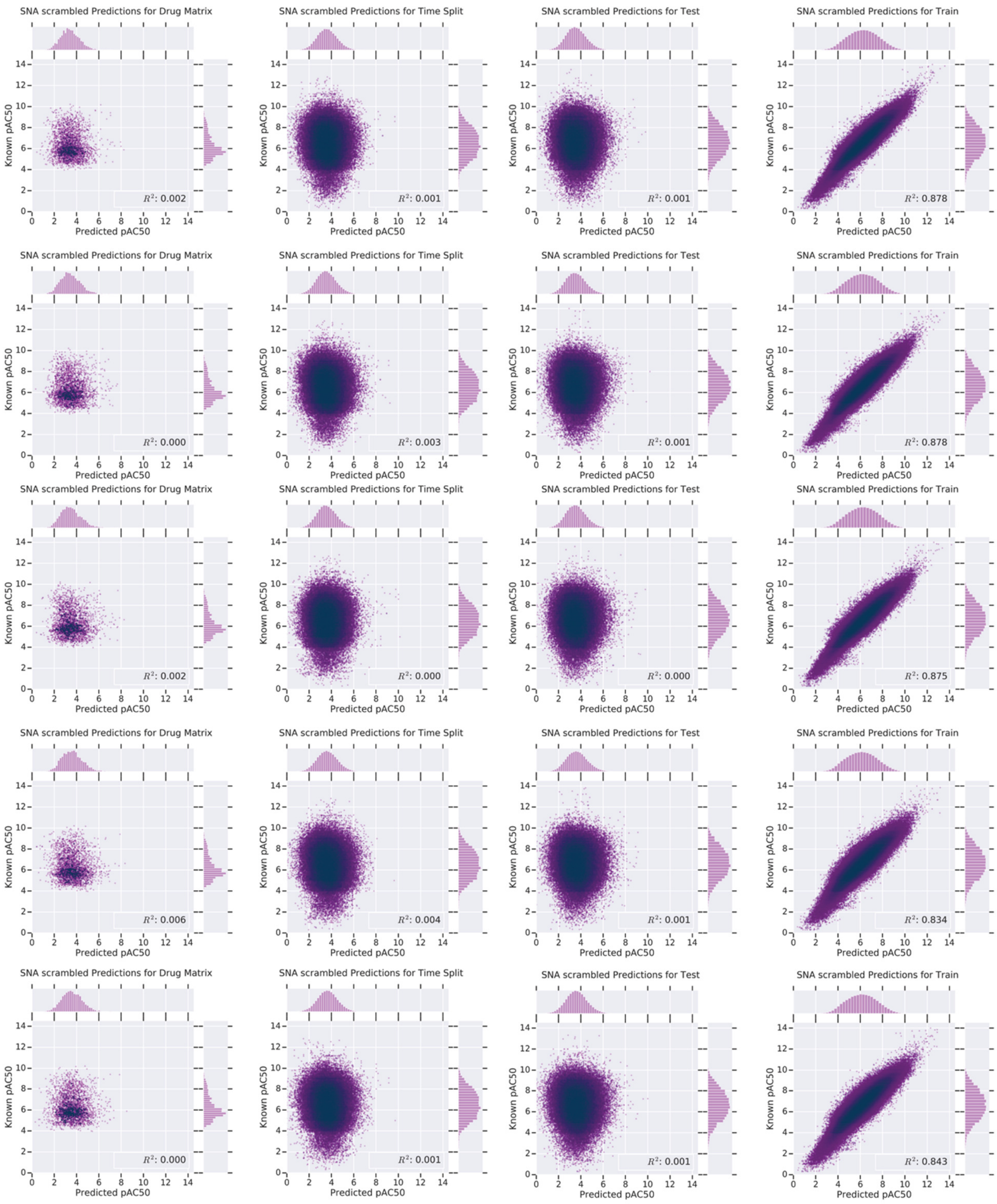

Supplementary Figure 14. *SNA scrambled* (y-randomized training set control with stochastic negatives) DNN $R^2$ plots for Drug Matrix (column 1), Time Split (column 2), Test (column 3), and Train (column 4) across each fold (0-4, top to bottom, increasing).

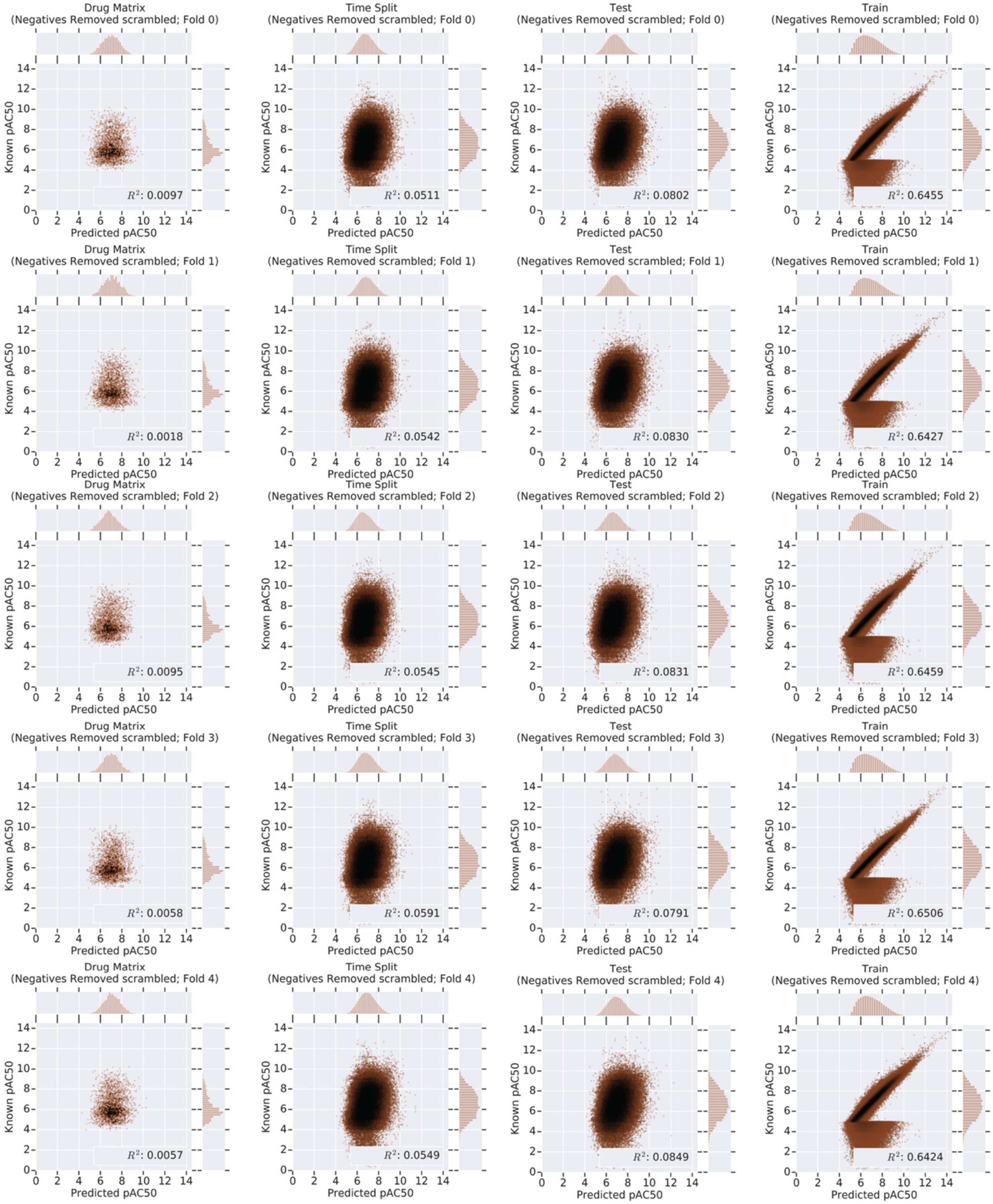

Supplementary Figure 15. *Negatives Removed scrambled* (y-randomized training set control with Negatives removed from the training set) DNN $R^2$ plots for Drug Matrix (column 1), Time Split (column 2), Test (column 3), and Train (column 4) across each fold (0-4, top to bottom, increasing).

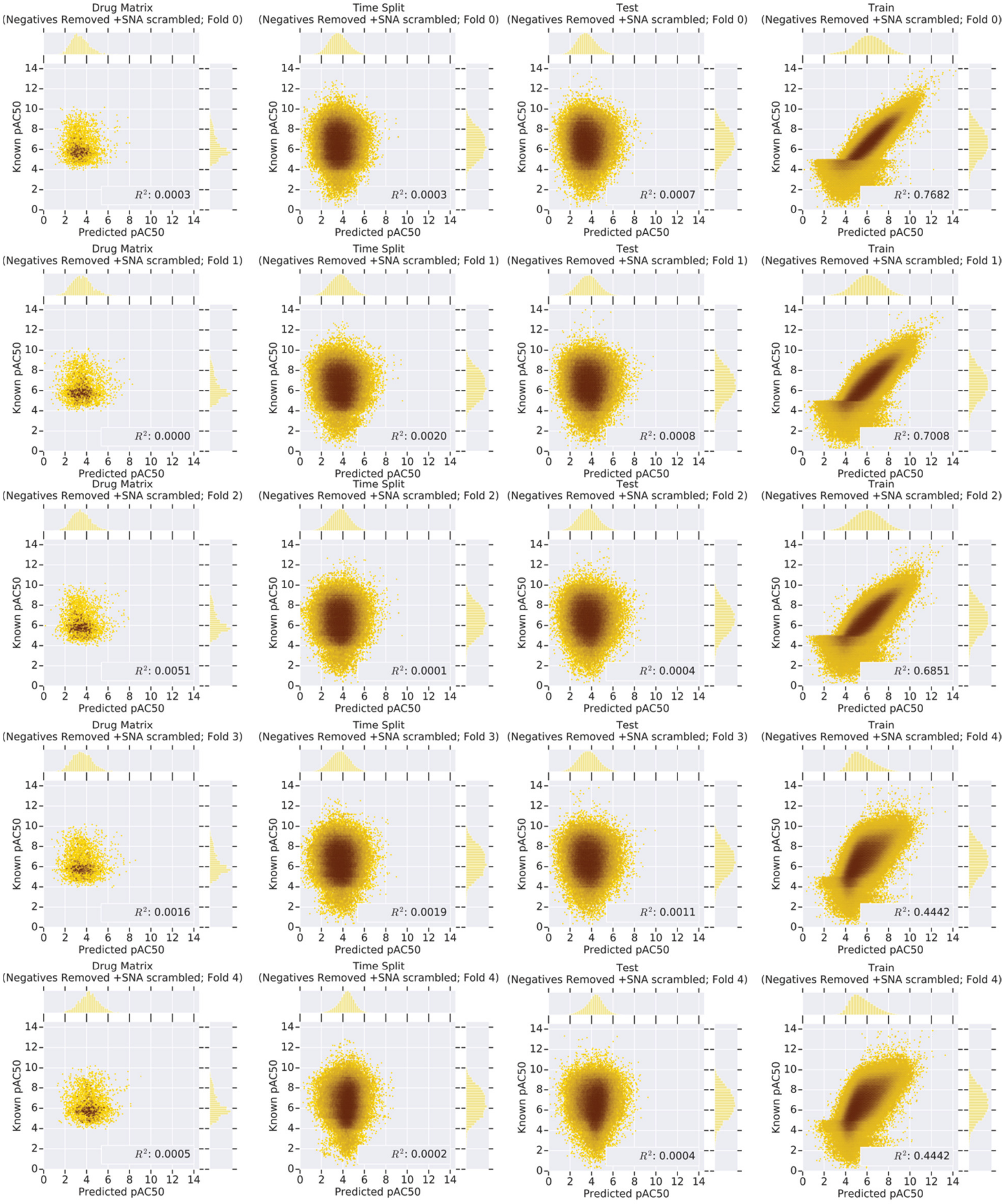

Supplementary Figure 16. *Negatives Removed +SNA scrambled* (y-randomized training set control with stochastic negatives) DNN $R^2$ plots for Drug Matrix (column 1), Time Split (column 2),  Test (column 3), and Train (column 4) across each fold (0-4, top to bottom, increasing).

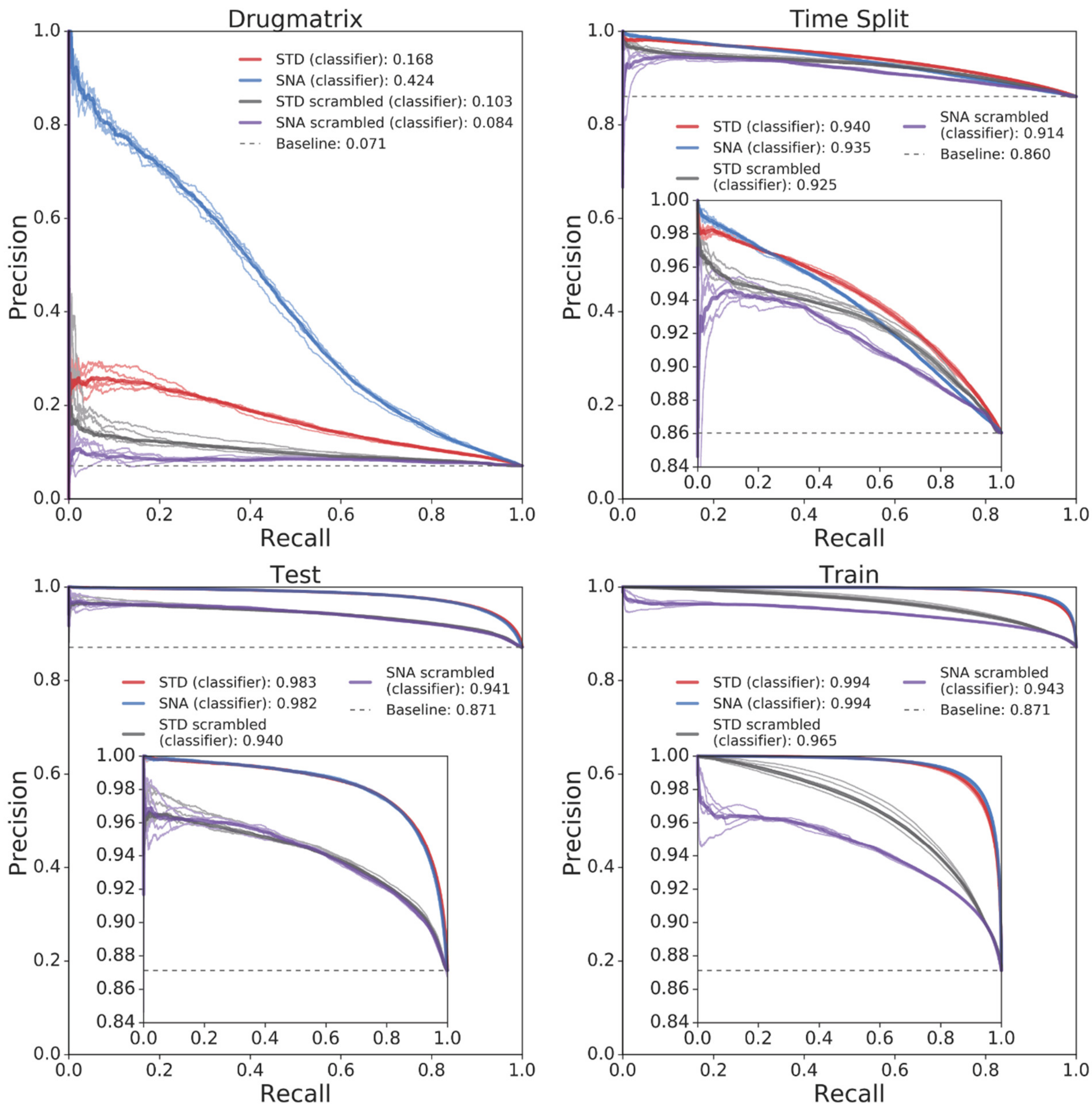

Supplementary Figure 17. AUPRC plots for *SNA, STD, SNA scrambled*, and *STD scrambled* classification DNNs for Drug Matrix (upper left), Time Split (upper right), Test (lower left), and Train (lower right). Each fold is plotted individually, with the mean AUPRC plotted with a thicker line.

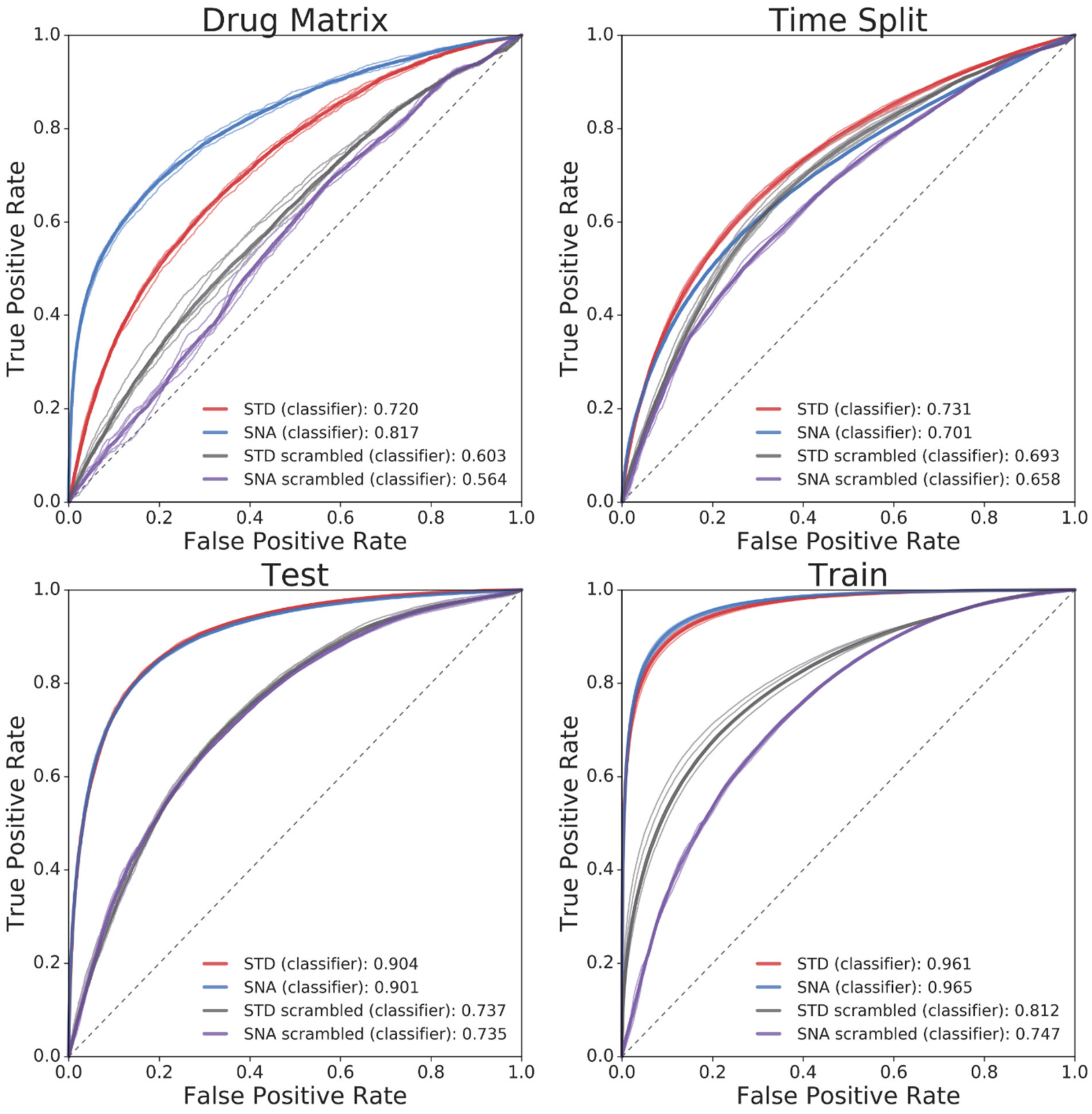

Supplementary Figure 18. AUROC plots for *SNA, STD, SNA scrambled*, and *STD scrambled* classification DNNs for Drug Matrix (upper left), Time Split (upper right), Test (lower left), and Train (lower right). Each fold is plotted individually, with the mean AUPRC plotted with a thicker line.

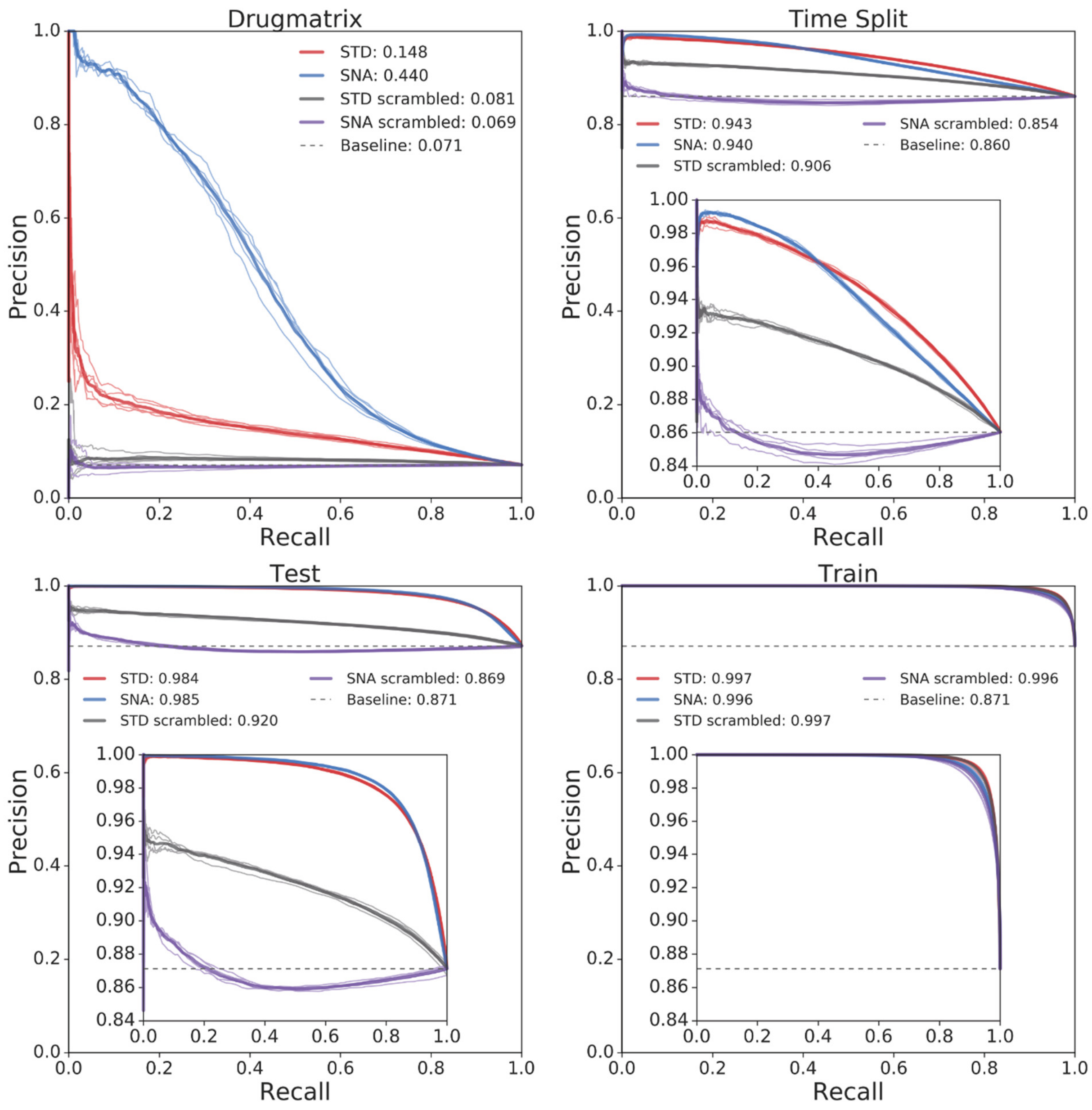

Supplementary Figure 19. AUPRC$_r$ plots for *SNA, STD, SNA scrambled*, and *STD scrambled* regression DNNs for Drug Matrix (upper left), Time Split (upper right), Test (lower left), and Train (lower right). Each fold is plotted individually, with the mean AUPRC plotted with a thicker line.

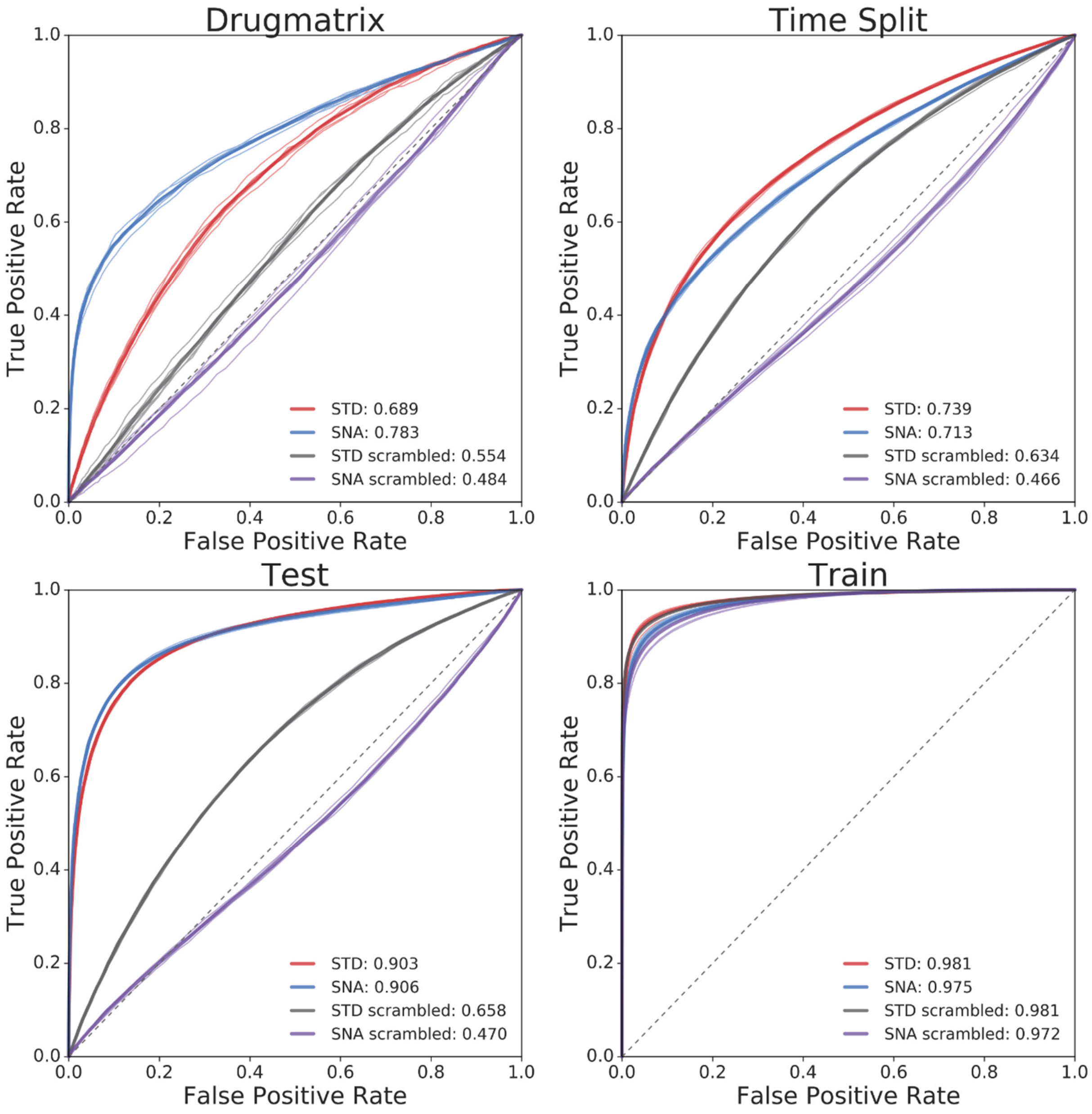

Supplementary Figure 20. $AUROC_r$ plots for *SNA, STD, SNA scrambled*, and *STD scrambled* regression DNNs for Drug Matrix (upper left), Time Split (upper right), Test (lower left), and Train (lower right). Each fold is plotted individually, with the mean AUPRC plotted with a thicker line.

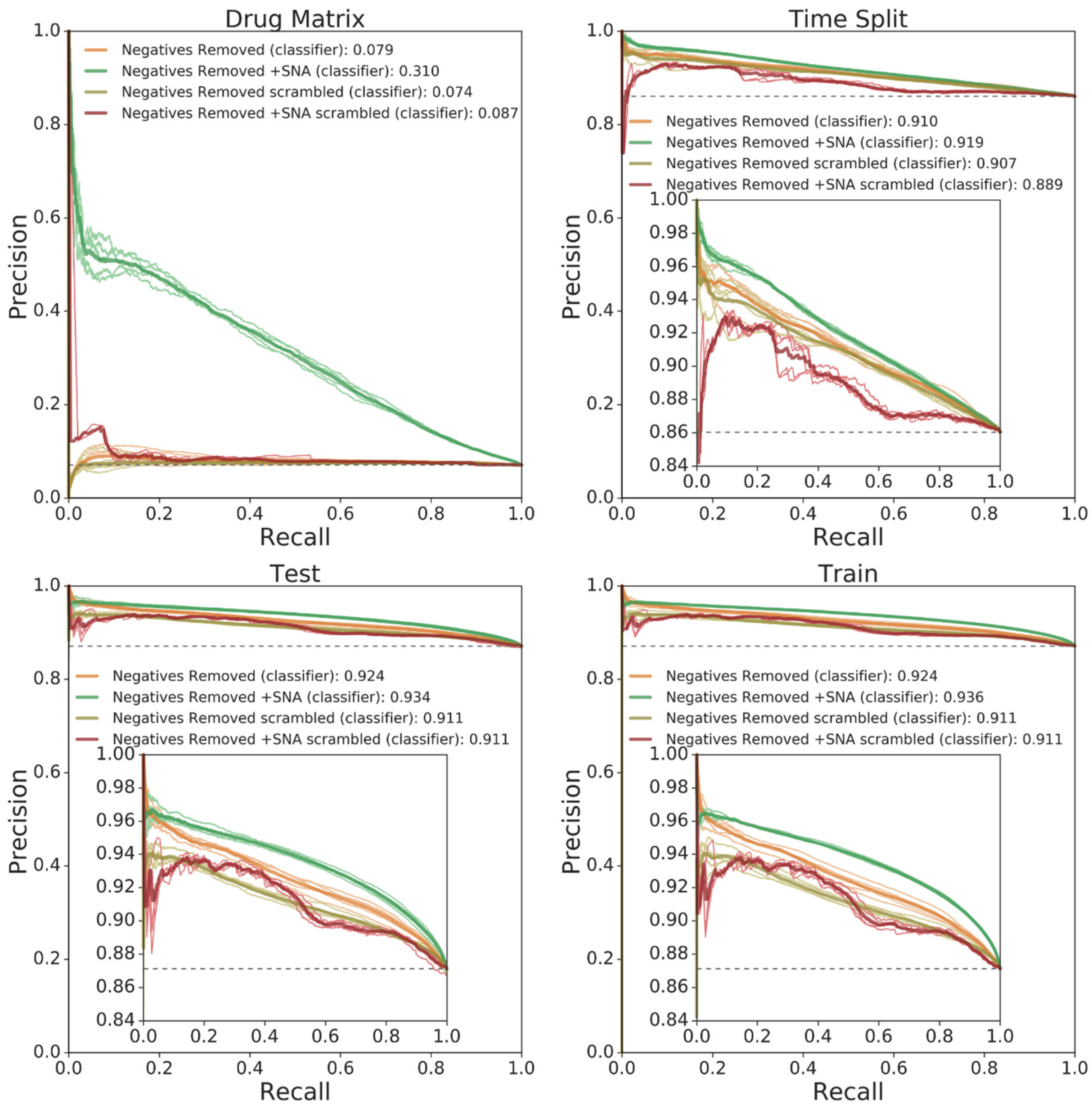

Supplementary Figure 21. AUPRC plots for *Negatives Removed, Negatives Removed +SNA, Negatives Removed scrambled*, and *Negatives Removed +SNA scrambled* classification DNNs for Drug Matrix (upper left), Time Split (upper right), Test (lower left), and Train (lower right). Each fold is plotted individually, with the mean AUPRC plotted with a thicker line.

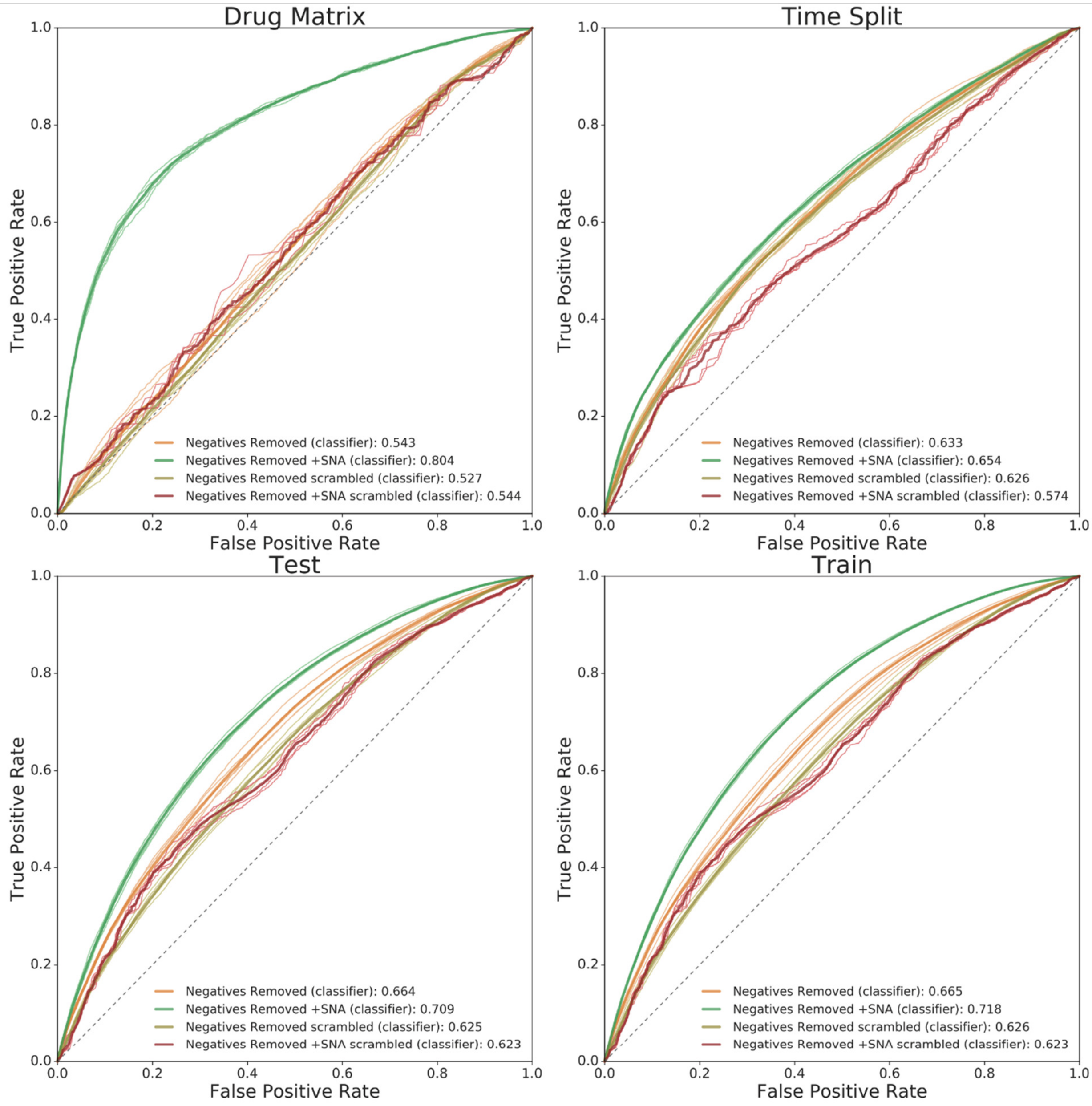

Supplementary Figure 22. AUROC plots for *Negatives Removed, Negatives Removed +SNA, Negatives Removed scrambled*, and *Negatives Removed +SNA scrambled* classification DNNs for Drug Matrix (upper left), Time Split (upper right), Test (lower left), and Train (lower right). Each fold is plotted individually, with the mean AUPRC plotted with a thicker line.

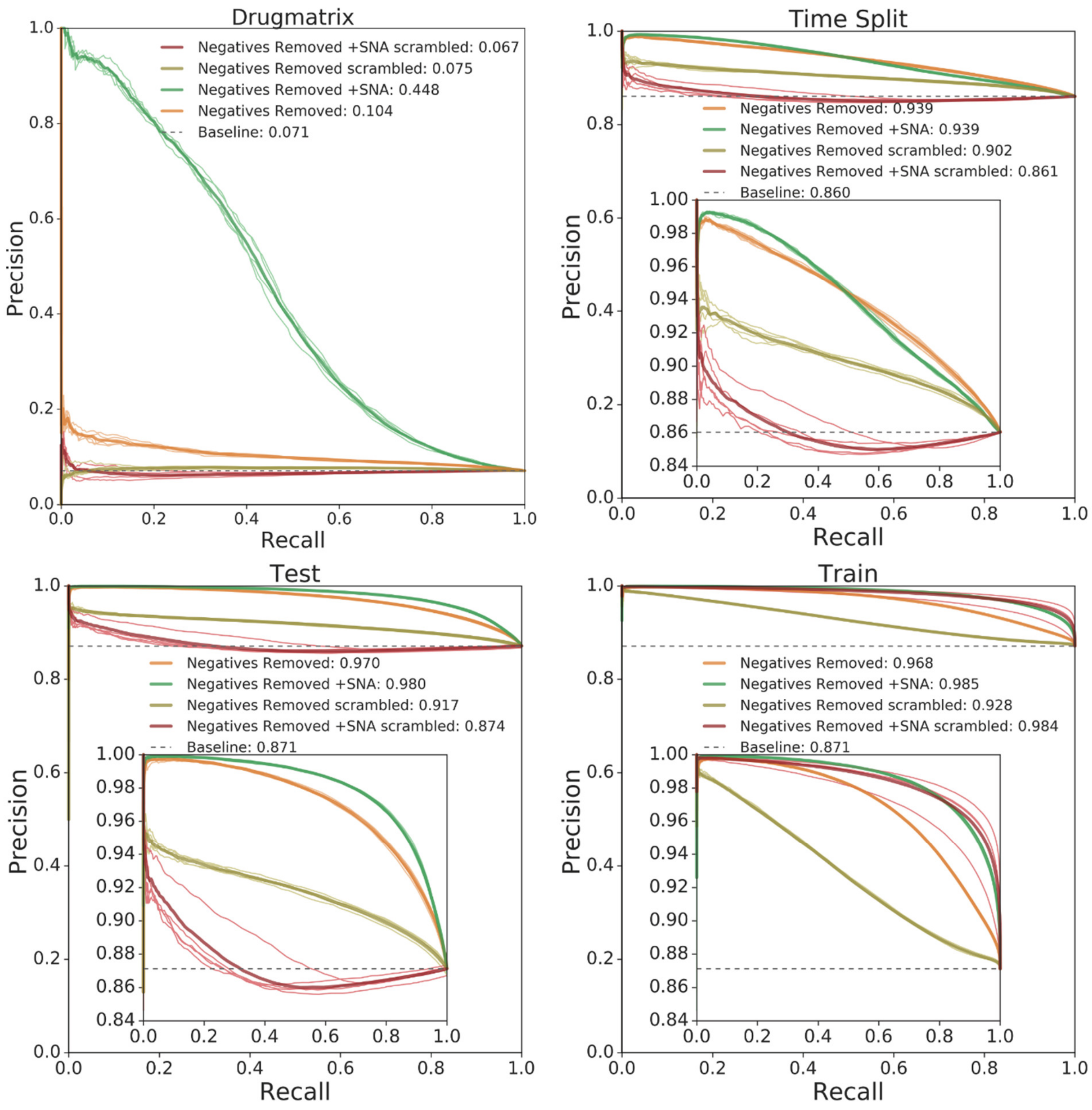

Supplementary Figure 23. AUPRC$_r$ plots for *Negatives Removed, Negatives Removed +SNA, Negatives Removed scrambled*, and *Negatives Removed +SNA scrambled* regression DNNs for Drug Matrix (upper left), Time Split (upper right), Test (lower left), and Train (lower right). Each fold is plotted individually, with the mean AUPRC plotted with a thicker line.

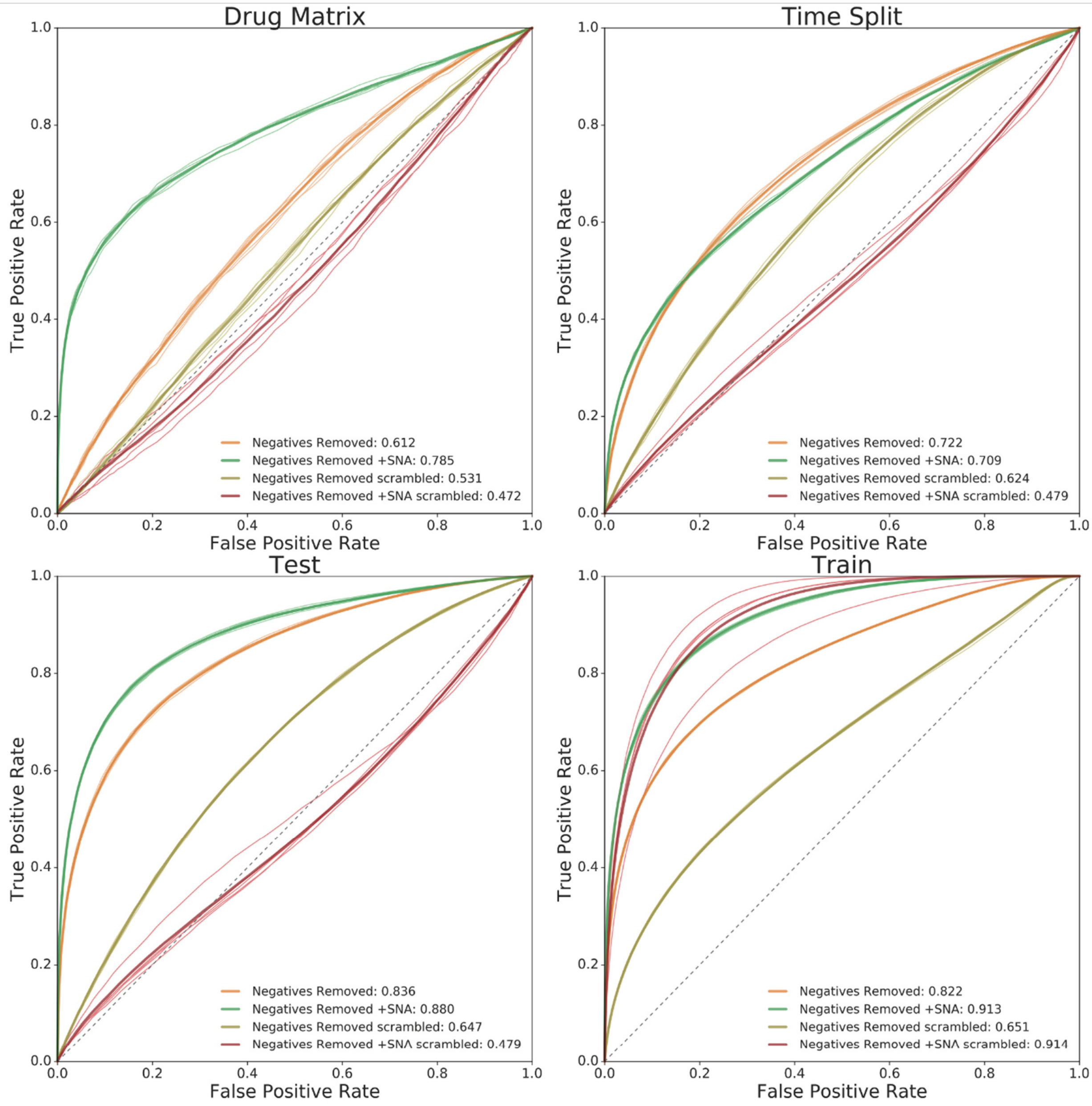

Supplementary Figure 24. AUPRC$_r$ plots for *Negatives Removed, Negatives Removed +SNA, Negatives Removed scrambled*, and *Negatives Removed +SNA scrambled* regression DNNs for Drug Matrix (upper left), Time Split (upper right), Test (lower left), and Train (lower right). Each fold is plotted individually, with the mean AUPRC plotted with a thicker line.
